# DeepSomatic: Accurate somatic small variant discovery for multiple sequencing technologies

**DOI:** 10.1101/2024.08.16.608331

**Authors:** Jimin Park, Daniel E. Cook, Pi-Chuan Chang, Alexey Kolesnikov, Lucas Brambrink, Juan Carlos Mier, Joshua Gardner, Brandy McNulty, Samuel Sacco, Ayse Keskus, Asher Bryant, Tanveer Ahmad, Jyoti Shetty, Yongmei Zhao, Bao Tran, Giuseppe Narzisi, Adrienne Helland, Byunggil Yoo, Irina Pushel, Lisa A. Lansdon, Chengpeng Bi, Adam Walter, Margaret Gibson, Tomi Pastinen, Midhat S. Farooqi, Nicolas Robine, Karen H. Miga, Andrew Carroll, Mikhail Kolmogorov, Benedict Paten, Kishwar Shafin

**Affiliations:** UC Santa Cruz Genomics Institute, University of California, Santa Cruz, CA, USA; Google Inc, Mountain View, CA, USA; Center for Cancer Research, National Cancer Institute, NIH, Bethesda, MD, USA; Sequencing Facility, Cancer Research Technology Program, Frederick National Laboratory for Cancer Research, Frederick, MD, USA; Sequencing Facility Bioinformatics Group, Biomedical Informatics and Data Science Directorate, Frederick National Laboratory for Cancer Research, Frederick, MD, USA; New York Genome Center, NY, USA; Children’s Mercy Hospital, University of Missouri-Kansas City School of Medicine, Kansas City, MO, USA

## Abstract

Somatic variant detection is an integral part of cancer genomics analysis. While most methods have focused on short-read sequencing, long-read technologies now offer potential advantages in terms of repeat mapping and variant phasing. We present DeepSomatic, a deep learning method for detecting somatic SNVs and insertions and deletions (indels) from both short-read and long-read data, with modes for whole-genome and exome sequencing, and able to run on tumor-normal, tumor-only, and with FFPE-prepared samples. To help address the dearth of publicly available training and benchmarking data for somatic variant detection, we generated and make openly available a dataset of five matched tumor-normal cell line pairs sequenced with Illumina, PacBio HiFi, and Oxford Nanopore Technologies, along with benchmark variant sets. Across samples and technologies (short-read and long-read), DeepSomatic consistently outperforms existing callers, particularly for indels.

## Introduction

Cancer is a genomic disease, with typical cancers containing thousands to tens of thousands of individual somatic variants distributed across a mosaic cell population of different clones (Stratton, Campbell, and Futreal 2009). The mutational processes that give rise to somatic variants are numerous and varied, with each cancer typically affected by a mixture of these processes over time (Alexandrov et al. 2020; Alexandrov and Stratton 2014). As a precise and accurate characterization of somatic variation reveals more of the pan-cancer mutational landscape, there has been an ongoing movement to bring somatic variant discovery into precision oncology and clinical practices (Perera-Bel et al. 2018; Garcia-Prieto et al. 2022; Farswan et al. 2021).

Somatic variation calling methods generally use tumor or combined tumor-normal sequencing to identify positions that differ from the reference genome, but are not germline variants. Most cancer genomics projects currently use short-read sequencing, which exhibits high base-level accuracy but is limited in its ability to map reliably to repetitive regions (W. Li and Freudenberg 2014). A multitude of short-read-based somatic variant callers have been developed, assessed, and applied to clinical research, with Strelka2 and MuTect2 being two top-performing tools across several previous benchmarking studies (Z. Chen et al. 2020; Pei et al. 2021; Garcia-Prieto et al. 2022; Jin et al. 2022; Farswan et al. 2021; Larson et al. 2012). Strelka2 uses a combination of mixture-model estimations for indel-calling, haplotype-modeling, and a normal sample contamination model, and exhibits faster run-time compared to other variant callers (Kim et al. 2018). MuTect2 is a Bayesian classifier-based model and exhibits high specificity, which is an advantage in identifying reliable subsets of low-frequency somatic variants (Cibulskis et al. 2013). Performance of these variant callers has shown to be good; evaluated against the HCC1395-HCC1395BL tumor-normal set of somatic variants published by the Somatic Mutation Working Group at the Sequencing Quality Control Phase II Consortium (SEQC2), Strelka2 achieves an F1-score of 0.9616 and MuTect2 achieves an F1-score of 0.9521 for single-nucleotide variants (SNVs) (Fang et al. 2021; Zheng et al. 2023).

In contrast to short-read sequencing, current long-read sequencing methods produce reads that are two to three orders of magnitude longer, but which have higher base-level error rates. For example, Oxford Nanopore Technologies’ (ONT) platform can sequence raw reads with lengths routinely in the 10s of kilobases (kb), ranging up to 100s of kb, with an accuracy of >99%, and can even achieve accuracies of 99.9% with duplex data (Damaraju, Miller, and Miller 2024; Logsdon, Vollger, and Eichler 2020; Kolesnikov et al. 2024). Similarly, Pacific Biosciences (PacBio)’s HiFi Revio system generates read lengths ranging from 10 to 25 kb, with accuracies of >99.9% (Damaraju, Miller, and Miller 2024). The accuracies of these long-read technologies are now comparable to Illumina short-read’s base-level accuracies of >99.9% (Logsdon, Vollger, and Eichler 2020). Long-read germline variant calling, with tools such as Clair3 (Zheng et al. 2022) and DeepVariant (Poplin et al. 2018), outperforms short-read methods in SNV identification (Shafin et al. 2021; Kolmogorov et al. 2023). In addition, long-reads, by virtue of being able to span multiple adjacent variants, are able to produce accurate, read-based phasing of variants, including rare variants, at megabase phase block scales (Kolmogorov et al. 2023).

To date, somatic variant discovery with long-reads has received much less attention. In this paper, we present DeepSomatic, a short-read and long-read somatic small variant caller, adapted from our DeepVariant (Kolesnikov et al. 2024; Shafin et al. 2021; Poplin et al. 2018) software, a deep learning-based germline variant calling method. Briefly, DeepVariant takes a set of aligned reads and a reference genome, generates a set of pileup images, identifies candidate variants, and uses a convolutional neural network (CNN) to classify each candidate variant’s genotype (Poplin et al. 2018). We developed DeepSomatic by heavily modifying DeepVariant, in particular, altering the pileup images to contain both tumor and normal aligned reads.

The existing somatic variant calling method ClairS, which was created by the developers of Clair3, can identify somatic variants from short and long reads (Zheng et al. 2023). While very promising, one notable limitation of ClairS is that it was trained using simulated data due to the lack of accurate somatic benchmarking sets available to use as a training set (Zheng et al. 2023). While simulations can be very useful, they are limited by their assumptions; in general, in order for a supervised learning method to usefully learn, accurate, representative benchmarking training sets are necessary. This has been a particular challenge in the somatic space. While for germline variant callers there are seven Genome-in-a-Bottle (GIAB) reference samples that each provide millions of reference germline variants to train on, the HCC1395-HCC1395BL tumor-normal cell line published by SEQC2 is currently the only publicly available set of high-quality somatic benchmark variants (Fang et al. 2021), and contains only tens of thousands of somatic variants. This is particularly unfortunate given the known heterogeneity of somatic mutation, with individual cancers exhibiting a mixture of different underlying mutational signatures (Alexandrov et al. 2020).

We address the lack of available training data sets by collecting and sequencing a set of five matching tumor-normal cell lines with Illumina, Pacific Biosciences (PacBio) HiFi, and Oxford Nanopore Technologies (ONT) sequencing technologies, including generating additional confirmatory data for the HCC1395-HCC1395BL tumor-normal cell line. For each sample, with exception of the HCC1395 sample, we utilized DNA derived from the same cell culture to ensure consistency in the mixture of sequenced mutations. From this data, we generated five sets of high confidence variants, each supported by multiple sequencing technologies. We used this data, in addition to the existing SEQC2 benchmark data for the HCC1395 sample, to train DeepSomatic. These publicly available benchmark sets combining information from both short- and long-read sequencing data should prove useful for future benchmarking and development efforts. We demonstrate that DeepSomatic outperforms existing somatic variant callers, providing a new tool for the accurate determination of these variants across different sequencing platforms. In a joint study, the generated cell line sequencing is used to develop a new somatic structural variation (SV) detection tool called Severus (Keskus et al. 2024).

## Results

### DeepSomatic, a new short- and long-read somatic variant caller

In **Figure 1a**, we describe the somatic variant identification process with DeepSomatic using tumor-normal sequencing data. DeepSomatic uses a similar framework to DeepVariant that has three main steps for variant identification:

- **make_examples**: In make_examples, DeepSomatic takes reads from the tumor sample and normal sample to create a tensor-like representation of the read features. The features like read base, read base quality, mapping quality etc. are represented as channels as shown in **Figure 1b**. This first step identifies potential somatic variants as candidates. Any candidate variant observed in the tumor sample that is not observed in the normal sample above a certain threshold is considered a somatic variant candidate. Each candidate is then represented in a manner where the reads from the normal sample appear on top and reads from the tumor sample appear at the bottom. This tensor-like representation is then passed to a convolutional neural network (CNN) for classification.
- **call_variants**: In the call_variants stage, DeepSomatic takes the tensor-like representation of each candidate and evaluates it with the CNN to classify if the candidate is a reference or sequencing error, germline variant or somatic variant. This stage replaces the genotyping task of DeepVariant to identify variants as somatic in DeepSomatic.
- **postprocess_variants**: After the classification, postprocess_variants takes the prediction output from call_variants and tags each candidate variant with the classification it got as somatic, germline or reference allele.

**Figure 1:**
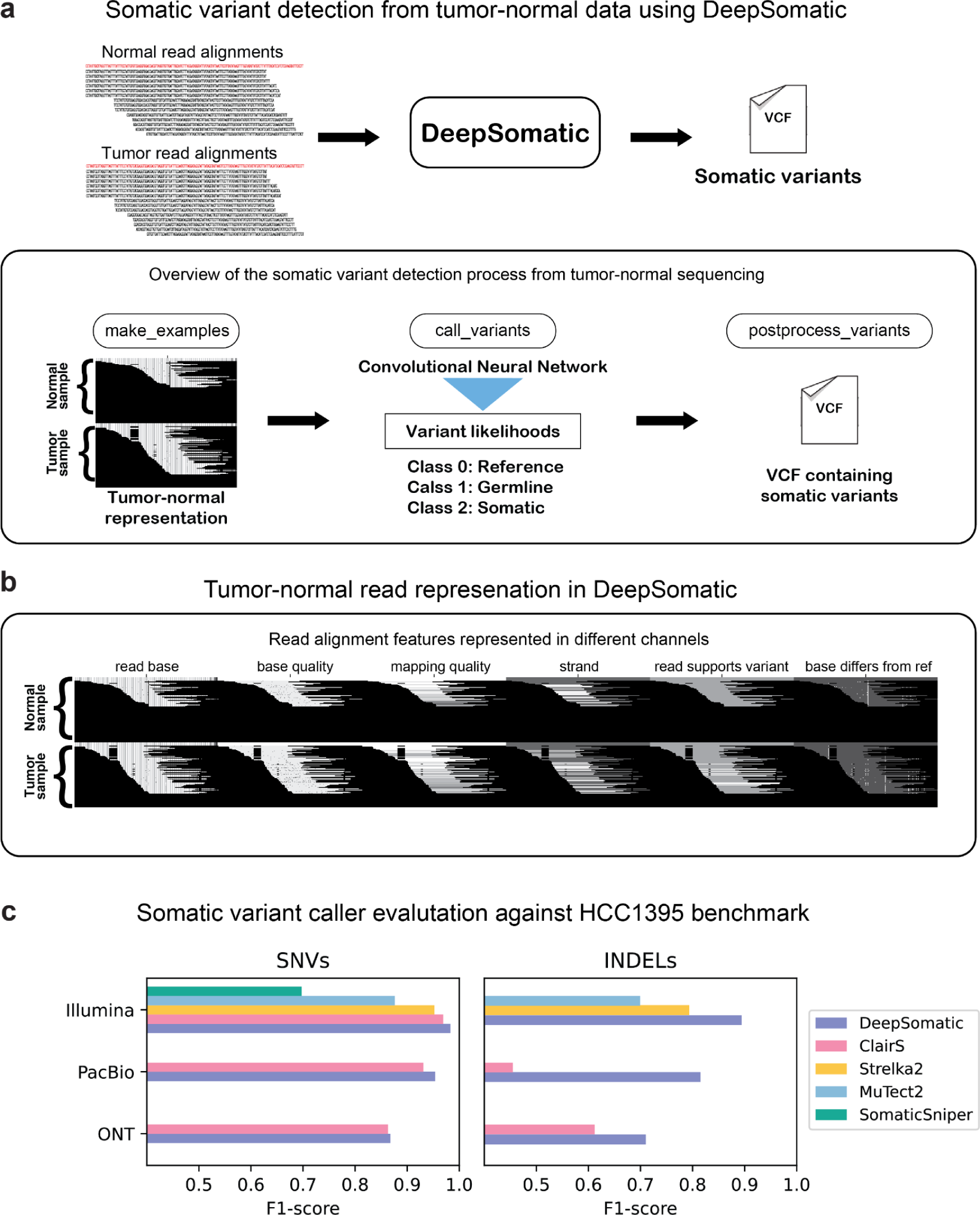
DeepSomatic overview and performance on SEQC2 HCC1395 benchmark. **(a)** Overview of DeepSomatic. **(b)** Tumor-normal read alignment features represented with six channels. **(c)** Somatic variant caller evaluation against SEQC2 HCC1395 benchmark for chromosome 1.

Training a deep-learning model like DeepSomatic requires a set of high-quality somatic training variants. The HCC1395-HCC1395BL tumor-normal breast cancer cell line, published by SEQC2, was the best available existing public benchmark for somatic small variants (Fang et. al 2021). We used this set of 41,072 somatic SNV and indel variants to train three initial sequence technology-specific models of DeepSomatic: HCC1395 Illumina, HCC1395 PacBio, and HCC1395 ONT.

For sequencing data, we used existing Illumina reads from SEQC2 (Fang et. al 2021) and Pacific Biosciences (PacBio) HiFi sequencing data from PacBio (https://www.pacb.com/connect/datasets), and additionally generated Oxford Nanopore Technologies (ONT) R10.4 chemistry sequencing data (**Supplementary Table 1**, methods). For training, using reads aligned to GRCh38, we use variants from chromosome 2 through chromosome 20 for training, and tune with chromosomes 21 and 22, while holding out chromosome 1 for validation. In all of the models we describe in this manuscript, we consistently use this training scheme of holding out chromosome 1 for validation. Overall, DeepSomatic models trained on HCC1395 data demonstrated improvements relative to existing variant callers for each sequencing technology (**Figure 1c**). Comparing Illumina models, the DeepSomatic HCC1395 model achieved the highest F1-scores of 0.9829 (SNVs) and 0.8944 (indels), while Strelka2 achieved F1-scores of 0.9521 (SNVs) and 0.7935 (indels), and ClairS achieved an F1-score of 0.9692 (SNVs) (**Figure 1c, Supplementary Table 2, 3**). Between PacBio models, the DeepSomatic HCC1395 model achieved F1-scores of 0.9536 (SNVs) and 0.8151 (indels) and ClairS achieved F1-scores of 0.9310 (SNVs) and 0.4550 (indels) (**Figure 1c, Supplementary Table 2, 3**). Lastly, between ONT models, the DeepSomatic HCC1395 model achieved F1-scores of 0.8677 (SNVs) and 0.7102 (indels) and ClairS achieved F1-scores of 0.8633 (SNVs) and 0.6122 (indels) (**Figure 1c**, **Supplementary Table 2, 3**).

### Sequencing five tumor-normal cell lines with multiple technologies

A variant caller that has been trained on only one sample can be limited in its ability to adapt to other samples, especially in the case of cancer data, because individual cancers demonstrate diverse mutational signatures (Alexandrov et al. 2020; Díaz-Gay et al. 2023). Therefore, a set of three breast cancer cell lines: HCC1395, HCC1937, HCC1954, and two lung cancer cell lines: H1437 and H2009 with matching normal cell lines were used to train new DeepSomatic models, which we call multi-cancer models, and generate sets of high-confidence somatic variants. We augmented the available HCC1395 data as discussed above; the remaining four cell lines were sequenced with Illumina, PacBio HiFi, and ONT R10.4 sequencing technologies, using DNA from the same cell culture extract for each of these cell lines to ensure consistency in the underlying set of variants assayed. Illumina reads were aligned using BWA-MEM2 (Vasimuddin et al. 2019), PacBio HiFi reads were aligned with pbmm2 (https://github.com/PacificBiosciences/pbmm2), and ONT reads were aligned with minimap2 (H. Li 2018). The PacBio alignments have about 60x tumor and 60x normal average coverage, and the ONT alignments have about 90x tumor and 30x normal coverage (**Figure 2a**). The Illumina alignments were downsampled to 100x tumor, 50x normal average coverage, in order to have consistent coverage with the long-read technologies. Long-read sequencing technologies have an N50 of about 18 kilobases (kb) for PacBio and 30-40 kb for ONT across the five cell lines (**Figure 2b**). The median read alignment identity to the GRCh38 reference for each sequencing technology is 0.9968 for PacBio, 0.9866 for ONT, and 1.0 for Illumina (**Figure 2c**). Additionally, datasets with various levels of tumor purity and normal purity were generated to mimic cases of tumor purity variance and tumor in normal (TIN) contamination that may occur in real tissue samples and included to train the DeepSomatic multi-cancer models. A summary of all datasets generated of the five tumor-normal cell lines for training is included in **Supplementary Table 1**. All data, including non-downsampled Illumina data, is publicly available for download without restriction (**Data Availability**).

**Figure 2:**
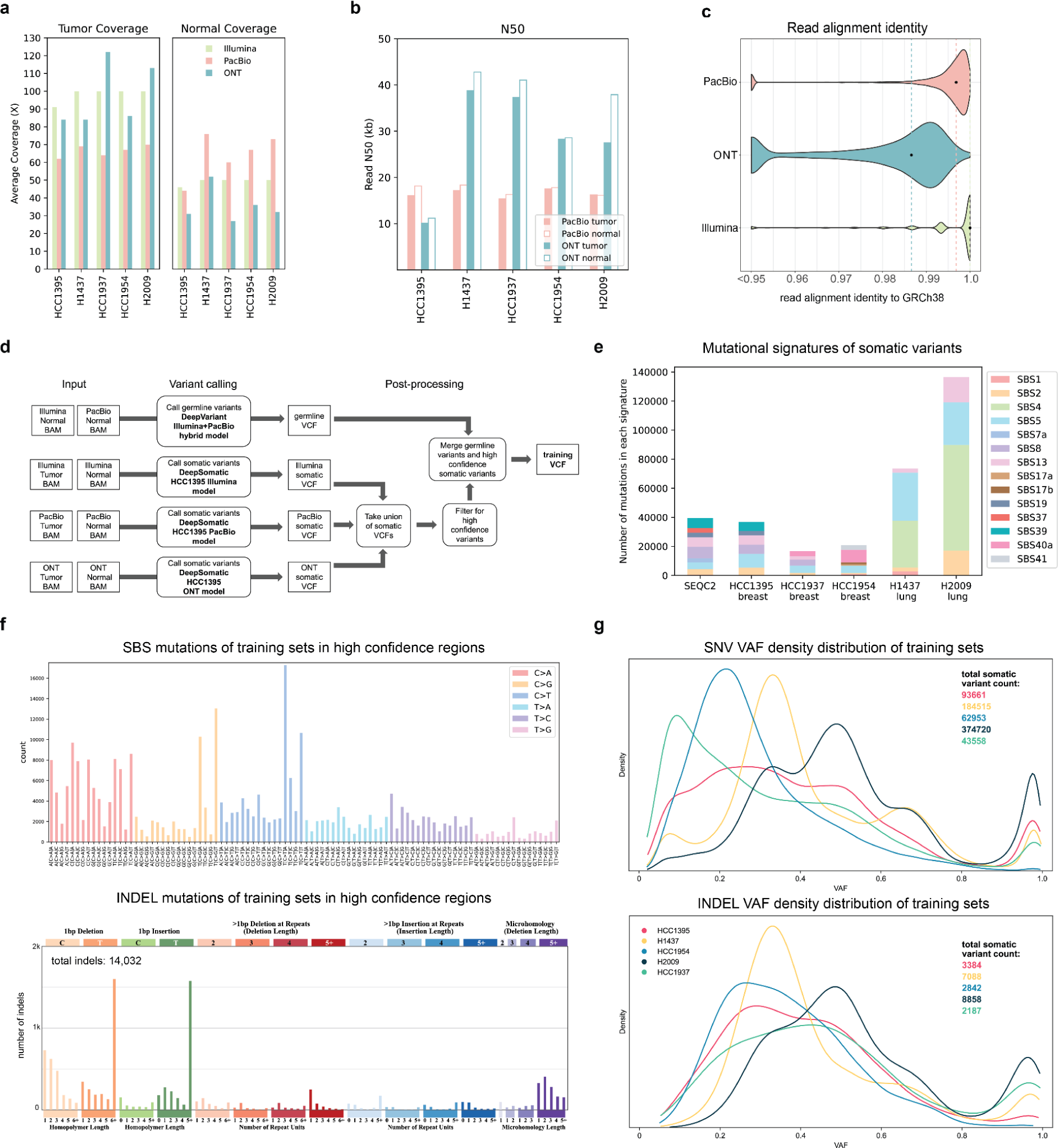
Training sets generated to train DeepSomatic. **(a)** Average sequencing fold coverage per sample. **(b)** Read N50 for long-read samples. **(c)** Read alignment identity to GRCh38 reference. Points mark the median for each sequencing technology. **(d)** Steps to derive training sets using DeepSomatic HCC1395 models. **(e)** Mutational signature analysis of the SEQC2 benchmark and somatic variants in training sets in high confidence regions derived as described in (d). **(f)** In high confidence regions aggregated across all five cell lines, (top) numbers of single base somatic substitutions stratified by context (SBS-96), and (below) types of indel mutations. Plots were generated using SigProfileMatrixGenerator (Bergstrom et al. 2019) **(g)** Variant allele frequency (VAF) distribution of somatic variants in training sets in high confidence regions represented as a Kernel Density Estimate (KDE) plot.

### Generating a high-quality benchmark set for five cell lines

For each tumor-normal cell line, the HCC1395 models were used to call somatic variants for each sequencing technology, to obtain a union of all somatic variants (**Figure 2d**). To create a subset of high-confidence variants for each line, the union set of all somatic variants was filtered using the criteria: 1) for SNVs, called by at least two out of the three HCC1395 models 2) for indels, called by the HCC1395 Illumina model and also called by at least one other long-read HCC1395 model. This filtering criteria requires that each variant included in the high-confidence set of variants is validated by at least two of the sequencing technologies and models.

High-quality germline variants were also called using DeepVariant’s Illumina+PacBio hybrid model and added to the final training set (**Figure 2d**). A set of high-confidence region BED files were also generated for each cell line by referencing methods used by SEQC2 to generate high-confidence regions for the published HCC1395 cell line (**Supplementary Figure 1, methods**). In addition, we subtracted SV regions called by Severus, segmental duplication (SD) regions defined by GIAB, and low-confidence variant regions that showed somatic variant signals, but did not have enough evidence to be considered a high-confidence somatic variant (**Supplementary Figure 1, methods**).

We verified our method for generating a subset of high-confidence variants by comparing the HCC1395 cell line’s high-confidence variants generated using our method to the existing SEQC2 benchmark set of variants. Our method calls 38,000 variants (93% of SEQC2) in common with the SEQC2 benchmark, and only calls 231 variants not present in the SEQC2 benchmark (**Supplementary Table 4**), while also showing highly similar mutational signatures (**Figure 2e**). Given their strong overlap, we opted to use the SEQC2 benchmark set for the HCC1395 cell line to be consistent with prior work.

Analysis of the training set of somatic variants for each cell line shows that each cell line has unique mutational characteristics, thus highlighting the importance of training our model with a diverse set of cell lines. A mutational signature analysis of the training set of somatic variants in high confidence regions for each cell line shows that different cancer types have different mutational signatures (**Figure 2e**). The number of different single base substitutions (SBS), analyzed using SigProfileMatrixGenerator (Bergstrom et al. 2019), shows that while, as expected, some mutations are much more prevalent (e.g. C>A mutations and C>T mutations), we capture hundreds of substitutions in every context (**Figure 2f**). SBS mutation analysis for each cell line depicts a distinct distribution amongst the different cell lines, illustrating, again, the unique mutational characteristics of each cell line (**Supplementary Figure 2**). Indel mutation analysis shows a notably larger number of T:A indels that are 5bp+ (**Figure 2f, Supplementary Figure 3**). A density distribution representation of VAFs (variant allele frequencies) for each of the training sets in high confidence regions also shows diverse VAF distributions reflecting the underlying aneuploidy and (potentially) subclonal structure of each cancer cell culture (**Figure 2g**).

### DeepSomatic performance on cell lines

Using the published set of somatic variants for HCC1395, and the training set somatic variants we generated for the other four cell lines, as well as including varying levels of tumor purities and TIN contamination for each of the cell lines, we trained three new DeepSomatic models: Illumina multi-cancer, PacBio multi-cancer, and ONT multi-cancer.

While the HCC1395 cell line has a published set of independent high-confidence somatic variants that can be used for evaluation, that is not the case for the other four cell lines. Therefore, to get around the circularity of evaluating our trained models using calls generated from an earlier generation of the same model, we evaluated the performance of our new DeepSomatic multi-cancer models using three different evaluation methods: 1) *chromosome 1 test set*: evaluating for chromosome 1, which was held out from training, using the benchmark sets we generated with the HCC1395 models; 2) *orthogonal tools benchmark*: a set we generated using a combination of the Strelka2 (Kim et. al 2018) and ClairS (Zheng et. al 2023) somatic variant callers; 3) *orthogonal technology benchmark*: evaluating for chromosome 1 (to, again, avoid circularity) against a benchmark set constructed by holding out the variant calls of the sequencing technology that is being evaluated.

The chromosome 1 test set evaluation directly assesses against the trained HCC1395 models; it is strongly biased toward the HCC1395 trained models because the set was derived directly from it, and is biased toward DeepSomatic in general. The orthogonal tools benchmark is a multi-sequencing technology benchmark, with the logic that a better variant caller is likely to agree with a consensus of existing methods. The orthogonal technology benchmark evaluation assesses the consistency of our individual technology models, assuming that accurate variant calls are likely to agree with the intersection of two independent sequencing technologies. None of these benchmarks is ideal, but together give indicators of overall performance.

Evaluating against the orthogonal technology and orthogonal tools benchmark sets shows that, overall, the DeepSomatic multi-cancer models have greater consistency amongst sequencing technologies than the DeepSomatic HCC1395 models as judged by overall higher F1-scores (**Figure 3a**, **Supplementary Figure 4**, **Supplementary Table 2**, **5**). The DeepSomatic multi-cancer models also outperform ClairS and Strelka2 for all five cell lines and three sequencing technologies when evaluated against the orthogonal technology benchmark (**Figure 3b**) and the chromosome 1 test set (**Supplementary Figure 5, Supplementary Table 3, 5**).

**Figure 3:**
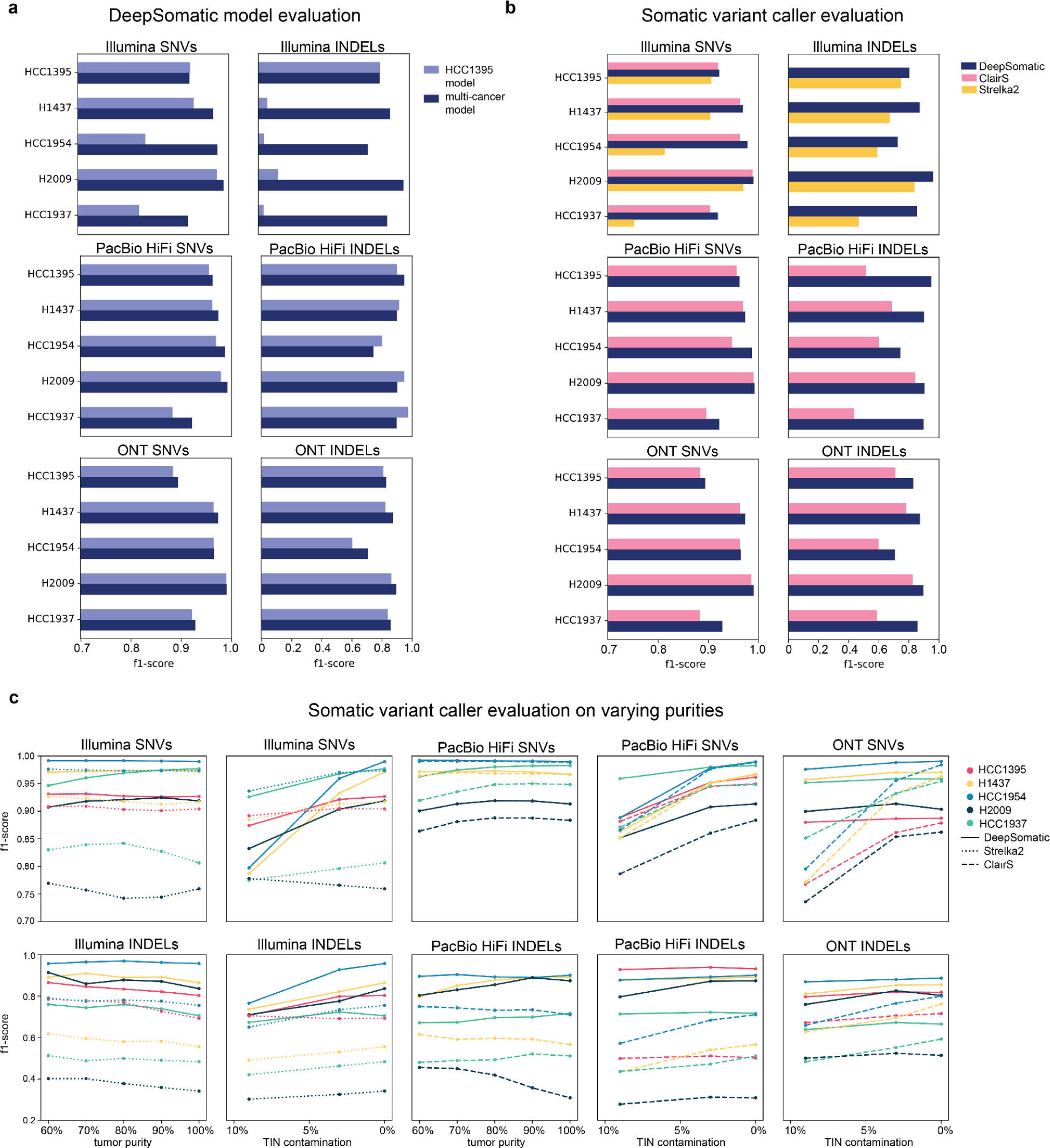
Somatic variant-calling performance of five tumor-normal cancer cell lines. Consistency evaluation is performed against the *orthogonal technology benchmark*. **(a)** Somatic variant calling f1-score performance of DeepSomatic HCC1395 model (trained with HCC1395 cell line only) and DeepSomatic multi-cancer model (trained with HCC1395 cell line plus four more tumor-normal cell lines). **(b)** Somatic variant calling f1-score performance of DeepSomatic multi-cancer model, ClairS, and Strelka2. **(c)** Somatic variant calling performance of DeepSomatic multi-cancer model, ClairS and Strelka2 on samples with varying purities.

Notably, using the orthogonal technology benchmark, the HCC1395 Illumina indels show much lower F1-scores in comparison to the multi-cancer model, due to a high number of false positives (**Figure 3a**, **Supplementary Table 2**). We investigated and found that these false positives are explained by differences in the way the reads were trimmed, something the multi-cancer model has learned to be robust to. This difference in Illumina reads is reflected in the lower read alignment identity of the four other cell lines compared to the HCC1395 cell line (**Supplementary Figure 6**).

The performance of the somatic variant callers on samples with varying levels of tumor purity was evaluated by running each model with varying tumor purities (90%, 80%, 70%, 60%) and 0% TIN contamination. And varying TIN contamination levels were evaluated (10% to 3%) with 100% tumor purity. These experiments had consistent coverage of 60x tumor and 35x normal. Overall, DeepSomatic models perform with consistent or increasing F1-scores with increasing tumor and normal purity, as expected (**Figure 3c, Supplementary Figure 7, Supplementary Table 6, 7, 8**). In comparison to Strelka2 and ClairS, DeepSomatic shows higher F1-scores across the different purity levels (**Figure 3c, Supplementary Figure 7, Supplementary Table 6, 7, 8**).

A high-confidence benchmark set of 304,663 somatic variants across five cell lines was generated using DeepSomatic’s multi-cancer models (**Supplementary Table 9**). Using this benchmark to evaluate Strelka2 and ClairS (it is circular to evaluate DeepSomatic here), we see that, overall, ClairS achieves higher F1-scores for SNVs, while Strelka2 achieves higher

F1-scores for indels (**Supplementary Figure 8, Supplementary Table 10**). Mutational signature analysis and variant allele frequency distributions of these benchmarks show high consistency to these analyses performed on the HCC1395 model derived benchmarks (**Figure 2e-g**, **Supplementary Figure 2, 3, 9, 10**).

### Extension of DeepSomatic to other types of cancer sequencing data

A matching normal sample is not always available due to additional sequencing costs or availability of the biological material. Instead, panels of normals - representing common germline variation in a population - can be used to distinguish somatic and germline variants (Cibulskis et al. 2013). To test this idea with DeepSomatic, three tumor-only DeepSomatic models, Illumina tumor-only, PacBio tumor-only, and ONT tumor-only, were trained using the HCC1395 benchmark from SEQC2 using a panel of normals (methods), with chromosome 1 held out. Evaluated against the HCC1395 benchmark on chromosome 1, these three models outperform ClairS across all technologies for both SNVs and indels (**Figure 4a, Supplementary Table 11**). Compared to corresponding tumor-normal models for both DeepSomatic and ClairS, we see that the tumor-only models achieve comparable recall values, especially for SNVs, but currently fall short in precision values (**Figure 4a**).

**Figure 4:**
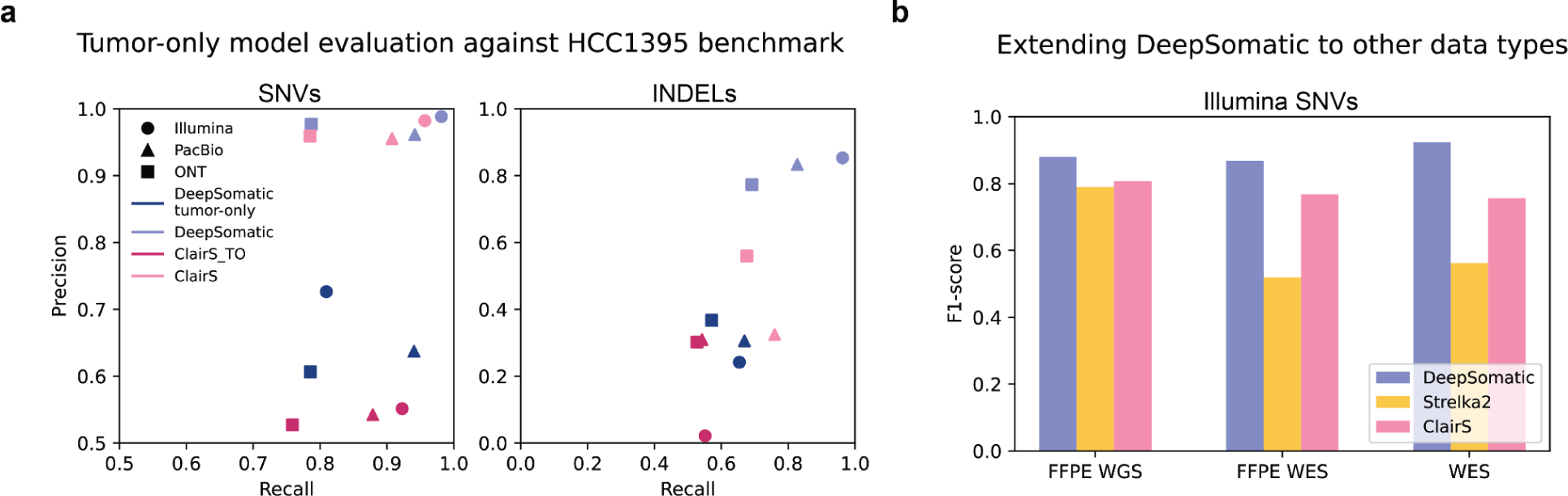
Extending DeepSomatic to tumor-only models and other data types. **(a)** Tumor-only somatic variant calling precision-recall performance of DeepSomatic tumor-only and ClairS_TO evaluated against SEQC2’s HCC1395 benchmark (chr1), with tumor-normal models included for reference **(b)** Somatic variant calling F1-score performance of DeepSomatic Illumina FFPE WGS, DeepSomatic Illumina FFPE WES and DeepSomatic Illumina WES models compared to Strelka2 and the ClairS Illumina model evaluated against SEQC2’s HCC1395 benchmark (chr1)

Clinical and archival samples are commonly preserved as FFPE (formalin-fixed paraffin-embedded) tissue samples. FFPE preservation typically introduces additional patterns of DNA damage (Steiert et al. 2023). DeepSomatic was extended to FFPE with two new models: FFPE_WGS and FFPE_WES. These models were trained on FFPE samples of the HCC1395 cell line published by SEQC2, and evaluated against the SEQC2 benchmarking set of variants on chromosome 1, which was excluded from training (Xiao et al. 2021; Fang et al. 2021). On FFPE WGS data, DeepSomatic’s FFPE_WGS model achieves a F1-score of 0.8803 (SNVs), while Strelka2 achieves a F1-score of 0.7894 (SNVs), and ClairS achieves a F1-score of 0.8072 (SNVs) (**Figure 4, Supplementary Table 11**). For indels, DeepSomatic’s FFPE_WGS model and Strelka2 achieved F1-scores of 0.8000 and 0.7894 (**Supplementary Table 11**).

WES (whole exome sequencing) is also commonly used because of cost-efficiency, but is limited to exomic regions and can also introduce sequencing artifacts (Koboldt 2020). DeepSomatic was also extended to WES cell line data in addition to the FFPE_WES model introduced above. The WES model was also trained on samples published by SEQC2 and both WES and FFPE_WES models were evaluated using the SEQC2 high confidence BED specific to exomic regions (Fang et al. 2021). On FFPE WES data, within a limited benchmark set of 152 SNV and indel variants, we report 45, 72 and 230 errors (FP+FN) for DeepSomatic’s FFPE_WES model, ClairS and Strelka2, respectively (**Figure 4, Supplementary Table 11**). On WES data, within the same benchmark set of 152 variants, we report 26, 82 and 218 errors for DeepSomatic’s FFPE_WES model, ClairS and Strelka2, respectively (**Figure 4, Supplementary Table 11**).

## Discussion

In this paper, we describe DeepSomatic, a highly accurate somatic small variant caller for Illumina short-reads, and PacBio HiFi and ONT long-reads. We also present a set of five matching tumor-normal cell lines with a supporting set of high-confidence somatic small variants for each cell line supported by multiple, independent sequencing technologies. We make all the underlying data and benchmarking sets publicly available without restriction to help alleviate the shortage of benchmarking resources for somatic variant calling. In a companion manuscript, we similarly investigate and generate novel sets of SVs for these lines (Keskus et al. 2024). Through a series of three evaluations across five cell lines, we conclude that DeepSomatic performs with higher F1-scores than existing short-read based somatic variant callers, and also achieves higher F1-scores than the single existing long-read based somatic variant caller, ClairS, especially for somatic indels.

Comparing Illumina, PacBio and ONT technologies for somatic variant calling, we see some evidence that PacBio long-reads improve accuracy relative to Illumina, with the average F1-score of our PacBio multi-cancer model across all five samples exceeding the Illumina model by 0.012 (SNVs) and 0.0354 (indels) using the orthogonal technology benchmark as measured by F1 balanced score. However, the currently higher read error rates of ONT do evidently reduce accuracy relative to PacBio, and Illumina, particularly for indels. We expect further chemistry and basecaller improvements to the ONT platform to rectify this.

The sequencing of an additional four tumor-normal lines increased the number of high-confidence benchmark variants by 263,591 variants (a 6.5 times increase) relative to the SEQC2 sets, and mutational signature analysis revealed quite different mutational patterns in the different cancers (Supplementary Table 9). For example, the two lung cancer cell lines, H1437 and H2009, show a large number of mutations in the SBS4 signature, which has a proposed etiology of tobacco smoking, that are not seen in the breast cancer cell lines (Alexandrov et. al 2020). The increased training data led to quantifiable improvements in benchmark performance, decreasing the total number of false positives (FP) and false negatives (FN) across the three technologies by 38.7% for SNVs and 95.2% for indels, with the part of significant decrease in indel errors attributed to the multicancer model learning to be robust to differences in read mapping differences. While this is gratifying, we believe that we’re likely quite far from reaching diminishing returns in terms of generating sufficient training data. Plausibly, the heterogeneity of cancer mutational signatures, the limited depth of sequencing we generated, the limitations of sequencing cell lines to represent cancer, and the fact that each cancer only contains order tens of thousands of mutations (in contrast to millions of germline variations used to train germline callers), are all good reasons to believe that the generation of additional, open sequencing data would be of significant benefit in creating open, accurate models. DeepSomatic can be easily re-trained using the additional sequencing data, suggesting it can be improved in the future as more cancer data benchmarks become available.

Beyond tumor-normal whole genome somatic variant calling, we created limited tumor-only, WES, and FFPE based modes of DeepSomatic. While it is ideal to have a matched tumor-normal pair sample, it is likely that there will be situations where only the tumor sample is available. Though still falling short of F1-scores of tumor-normal models, the tumor-only models achieve comparable recall values compared to the tumor-normal models, with recall values differing by about 0.02. DeepSomatic’s ability to be extended to other data types, such as FFPE data and WES data, shows that its structure is adaptable and applicable to relevant clinical experiments down the line. DeepSomatic’s models specific to FFPE and WES also prove to outperform other variant callers that were not specifically trained for these data types. The current limitation of FFPE and WES data for other cell lines, and low coverages for FFPE WES and WES data for the HCC1395 cell line that we evaluate on limits us from showing a more comprehensive performance evaluation for the FFPE and WES DeepSomatic models. This is important future work.

## Data Availability

Cell line sequencing produced in this study is openly available at NCBI SRA BioProject PRJNA1086849. Accession codes for the samples are organized in Supplementary Table 1.

Benchmarking sets derived from DeepSomatic and aligned reads used for training and evaluation are available in Google Cloud: [gs://brain-genomics-public/publications/park2024_deepsomatic](https://console.cloud.google.com/storage/browser/brain-genomics-public/publications/park2024_deepsomatic)

## Code Availability

DeepSomatic and DeepVariant code is available publicly on GitHub through (https://github.com/google/deepsomatic, https://github.com/google/deepvariant). Scripts for generating training and benchmarking sets are available on GitHub (https://github.com/jimin001/DeepSomatic_manuscript).

## Acknowledgements

HCC1395-HCC1395BL ONT sequencing was supported by the National Cancer Institute of the National Institutes of Health under Award Number U01CA253405. M.S.F. reports grants from Braden’s Hope for Childhood Cancer, Big Slick, Black & Veatch Foundation, Masonic Cancer Alliance, Noah’s Bandage Project, Elizabeth and Monte McDowell, Cancer Center Auxiliary, and the Department of Defense (W81XWH-20-1-0358). B.P. and J.P. were supported by National Institutes of Health (NIH) grants R01HG010485, U41HG010972, U24HG011853 and OT2OD033761. M.K. and A.K. were supported by the Intramural Research Program of the NIH.

## Author contributions

B.P., K.S, and M.K. helped conceive and direct the study. J.P. performed data analysis. D.E.C., P.C., A. Kolesnikov, L.B., J.C.M., A.C., and K.S. contributed to DeepSomatic development. J.P., A.B., and A. Keskus performed cell line data processing. J.G., B.M., and K.M. contributed to ONT data sequencing. S.S. performed Hi-C data sequencing. M.K., A. Keskus, A.B., and T.A. contributed to Severus development. J.S., Y.Z., and B.T. performed Illumina data sequencing. G.N., A.H., and N.R. performed ONT sequencing of the HCC1395 cell line. B.Y., I.P., L.A.L., C.B., A.W., M.G., T.P., and M.S.F. performed PacBio data sequencing. J.P., B.P., K.S., and M.K. drafted the manuscript.

## Methods

### Ethics statement

The Institutional Review Board of National Institutes of Health considers patient-derived cell lines as non-human subjects, and no approval was required. There are, however, ethical considerations, as the cell lines were derived prior to establishing the research use consent mechanism, and no such consent was received. Commercially available cell lines used in this study are anonymized, and the risks of identifying original patients or their immediate family members are low. On the other hand, openly releasing this data will significantly benefit research into developing new methods for detecting somatic variants - a critical task in current and future precision cancer therapies. Given the availability of existing data for these lines, we concluded that the benefits outweigh the marginal risks of the release of this additional data, and followed the practices established by the NCI and NHGRI in the TCGA tumor cell line data release (https://www.cancer.gov/ccg/research/genome-sequencing/tcga/history/ethics-policies).

### Multi-omics sequencing of five tumor/normal cell line pairs

Illumina, PacBio and ONT sequencing of five tumor/normal cell line pairs described in this study was generated jointly with a companion study (Keskus et al. 2024).

Cell lines were purchased from the American Type Culture Collection (ATCC). All cell lines were cultured according to ATCC handling guidelines at 37°C and 5% CO_2_. Cell counts were enumerated by Automated Cell Counter (BioRad, TC20), washed twice with Phosphate Buffered Saline (Gibco, 10010023), placed into 6 x 10^6^ cell aliquots, flash frozen in liquid nitrogen, and stored at -80°C. Cancer cell lines: HCC1954 (ATCC, CRL-2338), H2009 (ATCC, CRL-5911), HCC1937 (ATCC, CRL-2336), H1437 (ATCC, CRL-5872), and HCC1395 (ATCC, CRL-2324). Normal cell lines: HCC1954 BL (ATCC, CRL-2339), BL2009 (ATCC, CRL-5961), HCC1937BL (ATCC, CRL-2337), BL1437 (ATCC, CRL-5958), and HCC1395 BL (ATCC, CRL-2325). Utilizing the Monarch® HMW DNA Extraction Kit for Tissue (New England Biolabs, T3060), high molecular weight (HMW) DNA was extracted from 6 x 10^6^ cells for all cell lines.

### ONT sequencing of five tumor/normal cell line pairs

HMW DNA was sheared to a target size of 50kb using the Megaruptor 3 and DNAFluid+ kit (Diagenode, E07020001). DNA was quantified using the dsDNA BR assay on a Qubit fluorometer (ThermoFisher, Q33265), and size distribution was analyzed on Femto Pulse using the gDNA 165kb Analysis Kit (Agilent, FP-1002-0275). An SRE Kit (PacBio,102-208-300) was used to deplete DNA fragments <25kb in length. Libraries were created using an Ligation Sequencing Kit V14 (ONT, SQK-LSK114) following the standard kit protocol. Three library preps were conducted for cancer cell lines and one for normal cell lines. Cancer cell lines were sequenced on three flow cells, and normal cell lines were sequenced on one flow cell.

Libraries were eluted in 100µl of elution buffer. Sequencing was performed with version R10.4.1 PromethION flow cells (ONT, FLO-PRO114M) per manufacturer’s guidelines with the following alterations to maximize throughput; each library consisted of 25µL of DNA, 50µL of Load Beads, and 75µL of Sequencing Buffer to allow for four 150µl library loads per prep, and each flowcell was washed with the Flow Cell Wash Kit (ONT, EXP-WSH004) and reloaded every 24 hours, for a total runtime of 96 hours with four library loads.

### PacBio sequencing of five tumor/normal cell line pairs

Tumor and normal DNA was prepared for HiFi sequencing as previously described (Cohen et al. 2022) with the following modifications. Briefly, DNA was sheared to a target size of 14 kb using the Diagenode Megaruptor3, and SMRTbell libraries were prepared with the SMRTbell Express Template Prep Kit 3.0. Libraries were sequenced on Revio instrumentation (Pacific Biosciences, Menlo Park, CA) with 24 hr movies/SMRT cell. All samples were sequenced to a target depth of ∼60X coverage (2 SMRT cells per sample).

### Illumina sequencing of five tumor/normal cell line pairs

We used the TruSeq Nano DNA Prep from Illumina to prepare libraries. We took 200 ng of genomic DNA and fragmented it to a 400 bp insert size on a Covaris instrument which generates dsDNA fragments with 3’ or 5’ overhangs. The sheared DNA is blunt-ended and library size selection was performed using sample purification beads. A single ‘A’ nucleotide is added to the 3’ ends of the blunt fragments to prevent them from ligating to each other during the adapter ligation reaction. A corresponding single ‘T’ nucleotide on the 3’ end of the adapter provides a complementary overhang for ligating the adapter to the fragment. The indexed adapters are ligated to the ends of the DNA fragments and then PCR-amplified to enrich for fragments that have adapters on both ends. The final purified product is then quantitated by qPCR before cluster generation and paired-end sequencing (2 x150 bp) on the Illumina NovaSeq 6000 sequencer.

### ONT sequencing of HCC1395

The sample was fragmented to 20kb using Covaris g-TUBEs (COVARIS, 520079). DNA repair and end-prep was performed using the NEBNext Companion Module for Oxford Nanopore Technologies (NEB, E7180S) incubating at 20o C for 5 min followed by 5 min at 65o C. The sample was cleaned up using 1X Ampure XP beads (Beckman Coulter, A63881), washing twice on a magnetic rack with 80% ethanol. Sequencing adaptors and ligation buffer from the Oxford Nanopore Ligation Sequencing Kit (ONT, LSK1114) were ligated to DNA ends using Quick T4 DNA Ligase (NEB, E6056) for 45 min at room temp. The sample was cleaned up using 0.45X Ampure XP beads (Beckman Coulter, A63881), washing twice on a magnetic rack with the long-fragment buffer (ONT, LSK114) before eluting in 32uL of elution buffer (ONT, LSK114). Sequencing libraries were prepared by adding the following to the eluate: 100uL sequencing buffer (ONT, LSK114), 68uL loading solution (ONT, LSK114), and 0.5uL sequencing tether (ONT, LSK114). Samples were run on a PromethION (ver R10.4.1) flow cell using the PromethION sequencer. Sequencing runs were operated using the MinKNOW software (ver 22.12.5).

### Aligning five tumor-normal cell lines

All reads were aligned to the hg38 human genome reference.

### Illumina

Illumina reads were aligned with BWA-MEM2 (v2.2.1) using default parameters.

### PacBio

PacBio HiFi reads were aligned with pbmm2 with ‘--min-length 50’.

### ONT

ONT reads aligned with minimap2 with ‘--k 17 -y -K 5G --eqx’.

### Read alignment identity, N50, coverage

Read alignment identity and N50 were computed using wambam (https://github.com/nanoporegenomics/wambam) for each sequencing technology. The Illumina reads were reformatted to convert ‘M’ to ‘=’ and ‘X’ for CIGAR strings, using bbmap reformat.sh (https://github.com/BioInfoTools/BBMap/blob/master/sh/reformat.sh) to be suitable for wambam, which works with ‘X’ formatted CIGAR strings. Coverage was calculated using samtools (v1.13).

### Downsampling bams

Illumina bams were downsampled for 100x tumor, 50x normal coverage using samtools. Full coverage data are available at NCBI SRA BioProject PRJNA1086849.

### DeepSomatic tumor-only variant calling method

In **Supplementary Figure 11**, we show the tumor-only variant calling process with DeepSomatic. In absence of a matched normal sequencing, we use population allele frequencies as a feature for calling somatic variants accurately. After identifying the somatic variants, we then use a panel of normals to filter for somatic variants only.

- **make_examples**: In tumor-only mode, the make_examples stage takes the tumor sequencing read alignments and a VCF that includes population allele frequencies. For Illumina data, we use DeepVariant 1KG variant calls (Baid et al. 2020; “An Integrated Map of Genetic Variation from 1,092 Human Genomes” 2012) with population variants observed above 0.05. For PacBio and ONT, we use the Consortium of Long Read Sequencing Database (CoLoRSdb) (Lake and Sequencing (CoLoRS) 2024) data similarly filtered for variants observed with allele frequency higher than 0.05. We use DeepVariant’s ability to represent population allele frequencies as a channel (N.-C. Chen et al. 2023) to represent the population allele frequency as an extra channel in the tensor-like representation of the tumor-only data.
- **call_variants**: Call variants stage uses the examples generated for each candidate in the previous stage to classify if a variant is somatic, germline or reference allele.
- **postprocess_variants**: The postprocess_variants step takes the output of call_variants and a panel of normals VCF to identify somatic-only variants. For short reads, a panel of normals is done using VCFs available from dbSNP (Sherry et al. 2001), gnomad (Karczewski et al. 2020) and 1000 genomes project (Auton et al. 2015). For long reads, we use the Consortium of Long Read Sequencing Database (CoLoRSdb) (Lake and Sequencing (CoLoRS) 2024) VCF with dbSNP and gnomad.

In **Supplementary Figure 11b**, we show all features represented in DeepSomatic tumor-only mode. In tumor-only mode, we have one extra channel to represent the population allele frequency with all the other features derived from the tumor-only sequencing like base, base-quality, mapping quality etc.

### DeepSomatic evaluation methods

#### Benchmarking

All VCFs were evaluated using som.py using docker image (pkrusche/hap.py:latest) with ‘-N --restrict-regions ${BED} --feature-table generic’.

### Somatic variant caller evaluation on SEQC2 benchmark

Somatic variant caller evaluation was performed using WGS_NS_N_1.bwa.dedup.bam, WGS_NS_T_1.bwa.dedup.bam, and WGS_NS_T_2.bwa.dedup.bam from SEQC2 (https://ftp-trace.ncbi.nlm.nih.gov/ReferenceSamples/seqc/Somatic_Mutation_WG/data/WGS/)

DeepSomatic was run using the HCC1395 models using default parameters. Strelka2 (v2.9.10) and ClairS (v0.2.0) were run using default parameters. Variant calls for MuTect2 and SomaticSniper are from SEQC2 (https://ftp-trace.ncbi.nlm.nih.gov/ReferenceSamples/seqc/Somatic_Mutation_WG/analysis/SNVs/vcfs/WGS/). Specific variant call sets used: WGS_NS_1.bwa.muTect2.vcf.gz, WGS_NS_1.bwa.somaticSniper.vcf.gz.

### Generating benchmarking set VCF files and high confidence region BED files using DeepSomatic HCC1395 models

#### VCF files

The training set VCFs used for training the DeepSomatic multi-cancer models were generated by taking a union set of all somatic variants from three DeepSomatic HCC1395 models for each sequencing technology. The union set was then filtered for high-confidence variants against a criteria of: 1) for SNVs, called by at least two out of the three HCC1395 models 2) for indels, called by the HCC1395 Illumina model and also called by at least one other HCC1395 model. This filtering criteria requires that each variant included in the high-confidence set of variants is validated by at least two of the sequencing technologies and models. High-quality germline variants were also called using DeepVariant’s Illumina+PacBio hybrid model and added to the final training set (**Figure 2a**).

#### BED files

Steps to generate high confidence region BED files were adapted from the methods used by SEQC2 in generating their HCC1395 benchmarking set (**Supplementary Figure 1**). For each cell line across all three sequencing technologies, GATK (broadinstitute/gatk3:3.8-1) CallalbeLoci was used to identify the callable regions for both tumor and normal bams. Parameters were set to ‘--maxDepth [8*average_coverage] --maxFractionOfReadsWithLowMAPQ 0.1 --maxLowMAPQ 1 --minMappingQuality 20 --minDepth 10 --minDepthForLowMAPQ 10’. ‘--minBaseQuality’ was set to 20 for Illumina and PacBio HiFi reads, and to 7 for ONT reads. Additional flags were also added: ‘--filter_reads_with_N_cigar --filter_mismatching_base_and_quals --filter_bases_not_stored --allow_potentially_misencoded_quality_scores’. The callable regions in both tumor and normal aligned reads for each sequencing technology were merged together as a union set of all callable regions. A series of three filtering steps were performed to remove these regions: 1) “Low confidence variant regions” identified by subtracting the high confidence variants included in the training sets from the union set of variants, 2) structural variant (SV) regions identified by Severus (+/- 1000bp removed from start/end of each SV), 3) segmental duplication regions published by GIAB. Additionally, loss of heterozygosity regions in normal bams, identified by Wakhan (https://github.com/KolmogorovLab/Wakhan), were removed from HCC1395 and H2009 cell lines. The percentage of the whole genome covered by these BED files are reported in **Supplementary Table 12**.

### Generating benchmarking VCF files and BED files

#### Orthogonal tools benchmark VCF files

The orthogonal tools benchmark was generated by taking a union of somatic variants from Strelka2, ClairS PacBio and ClairS ONT models, then filtering for high confidence variants against a criteria of: 1) for SNVs, called by at least two out of the three models 2) for indels, called by Strelka2 (Illumina) and also called by at least one of the ClairS models (long reads). The same orthogonal tools benchmark was used to evaluate both DeepSomatic HCC1395 and DeepSomatic multi-cancer models.

#### Orthogonal tools benchmark BED files

High confidence regions to evaluate alongside the orthogonal tools benchmark VCFs were generated using the GIAB HG002 BED file as a prior, and subtracting out “low confidence variant regions”. “Low confidence variant regions” were defined by subtracting the variant regions included in the orthogonal tools benchmarking VCF from the union set of somatic variants called by Strelka2, ClairS PacBio and ClairS ONT models.

#### Orthogonal technology benchmark VCF files

The orthogonal technology benchmark consists of a two-technology intersection set of multi-cancer PacBio and multi-cancer ONT model somatic calls to evaluate the multi-cancer Illumina model, an intersection set of multi-cancer Illumina and multi-cancer ONT model somatic calls to evaluate the multi-cancer PacBio model, and an intersection set of multi-cancer Illumina and multi-cancer PacBio somatic calls to evaluate the multi-cancer ONT model. A parallel version of these orthogonal technology benchmark sets were also generated for the HCC1395 models to evaluate the HCC1395 models.

#### Orthogonal technology benchmark BED files

High confidence regions to evaluate alongside the orthogonal technology benchmark VCFs were generated by using “callable regions” called by GATK CallableLoci, for the two technologies included in the VCF, then subtracting out “low confidence variant regions”, structural variant regions and segmental duplication regions.

### Generating benchmarking set VCF files and high confidence region BED files using DeepSomatic multi-cancer models

#### VCF files

A set of high confidence variants using the multi-cancer models was generated using the same method that was used for generating a high confidence set using the HCC1395 models. A union set was generated with all somatic variants from the three DeepSomatic multi-cancer models for each sequencing technology. The union set was then filtered for high-confidence variants against a criteria of: 1) for SNVs, called by at least two out of the three multi-cancer models 2) for indels, called by the multi-cancer Illumina model and also called by at least one other multi-cancer model. High-quality germline variants were also called using DeepVariant’s Illumina+PacBio hybrid model and added to the final training set.

#### BED files

High confidence regions for the multi-cancer models were defined using the same method that was used to generate high confidence regions for the HCC1395 models described above. The only difference between these high confidence regions are the “low confidence variant regions”, which were generated using variants called by the multi-cancer models instead of the HCC1395 models.

### Generating titration bam files

Titration bams were generated using samtools (v1.13). A subset of reads from each tumor bam and normal bam were extracted to set aside as reads that would be spiked in to simulate contamination, as to not cause duplicate reads. Random subsampling of reads within a bam was performed using ‘samtools view’ with ‘-s [seed + fraction to subsample]’. A different seed was used for subsampling the same bam file multiple times. More details on these steps are described in the command lines section and the scripts used to run these commands. For the HCC1395 cell line, titration bams published by the SEQC2 consortium were used for training.

### Mutational signature analysis

Mutational signature analysis was performed on high confidence variant sets generated using the DeepSomatic HCC1395 models and the DeepSomatic multi-cancer models using SigProfilerAssignment (https://github.com/AlexandrovLab/SigProfilerAssignment). The cosmic_fit function was used with cosmic_version 3.4 and context_type 96 using the hg38 reference genome.

### Mutational matrix generation

Mutational matrixes for SBS-96 and indels were created from the high confidence variant sets generated using the DeepSomatic HCC1395 models and the DeepSomatic multi-cancer models using SigProfilerMatrixGenerator (https://github.com/AlexandrovLab/SigProfilerMatrixGenerator). The SigProfilerMatrixGeneratorFunc function was used using the hg38 reference genome. Plots in Figure 2f were generated by SigProfilerMatrixGenerator.

## Supplementary

### Supplementary tables

Supplementary Table 1: [Sequencing data details and accessions](https://docs.google.com/spreadsheets/d/1TqJbOjNA5dS2Gwgm074IKo2gyKqzAqq-t6DtCFknghE/edit?gid=1192714176#gid=1192714176)

Supplementary Table 2: [DeepSomatic HCC1395 model evaluation statistics](https://docs.google.com/spreadsheets/d/1TqJbOjNA5dS2Gwgm074IKo2gyKqzAqq-t6DtCFknghE/edit?gid=1496791747#gid=1496791747)

Supplementary Table 3: [Other variant callers evaluation statistics](https://docs.google.com/spreadsheets/d/1TqJbOjNA5dS2Gwgm074IKo2gyKqzAqq-t6DtCFknghE/edit?gid=1944311675#gid=1944311675)

Supplementary Table 4. [Comparing variant counts in DeepSomatic derived benchmarks and SEQC2 benchmark in high confidence regions](https://docs.google.com/spreadsheets/d/1TqJbOjNA5dS2Gwgm074IKo2gyKqzAqq-t6DtCFknghE/edit?gid=278142881#gid=278142881)

Supplementary Table 5: [DeepSomatic multi-cancer model evaluation statistics](https://docs.google.com/spreadsheets/d/1TqJbOjNA5dS2Gwgm074IKo2gyKqzAqq-t6DtCFknghE/edit?gid=486338583#gid=486338583)

Supplementary Table 6: [Illumina purity evaluation of DeepSomatic, ClairS and Strelka2](https://docs.google.com/spreadsheets/d/1TqJbOjNA5dS2Gwgm074IKo2gyKqzAqq-t6DtCFknghE/edit?gid=1662432188#gid=1662432188)

Supplementary Table 7: [PacBio purity evaluation of DeepSomatic and ClairS](https://docs.google.com/spreadsheets/d/1TqJbOjNA5dS2Gwgm074IKo2gyKqzAqq-t6DtCFknghE/edit?gid=123403587#gid=123403587)

Supplementary Table 8: [ONT purity evaluation of DeepSomatic and ClairS](https://docs.google.com/spreadsheets/d/1TqJbOjNA5dS2Gwgm074IKo2gyKqzAqq-t6DtCFknghE/edit?gid=1697534283#gid=1697534283)

Supplementary Table 9: [Variant count in DeepSomatic derived benchmarking sets](https://docs.google.com/spreadsheets/d/1TqJbOjNA5dS2Gwgm074IKo2gyKqzAqq-t6DtCFknghE/edit?gid=1585344457#gid=1585344457)

Supplementary Table 10: [DeepSomatic multi-cancer model derived benchmarks to evaluate existing somatic variant callers](https://docs.google.com/spreadsheets/d/1TqJbOjNA5dS2Gwgm074IKo2gyKqzAqq-t6DtCFknghE/edit?gid=2102224829#gid=2102224829)

Supplementary Table 11: [Evaluation of FFPE, WES and tumor-only data types](https://docs.google.com/spreadsheets/d/1TqJbOjNA5dS2Gwgm074IKo2gyKqzAqq-t6DtCFknghE/edit?gid=1158416729#gid=1158416729)

Supplementary Table 12: [Percent of whole genome covered by BED files](https://docs.google.com/spreadsheets/d/1TqJbOjNA5dS2Gwgm074IKo2gyKqzAqq-t6DtCFknghE/edit?gid=41901748#gid=41901748)

### Supplementary figures

**Supplementary Figure 1:**
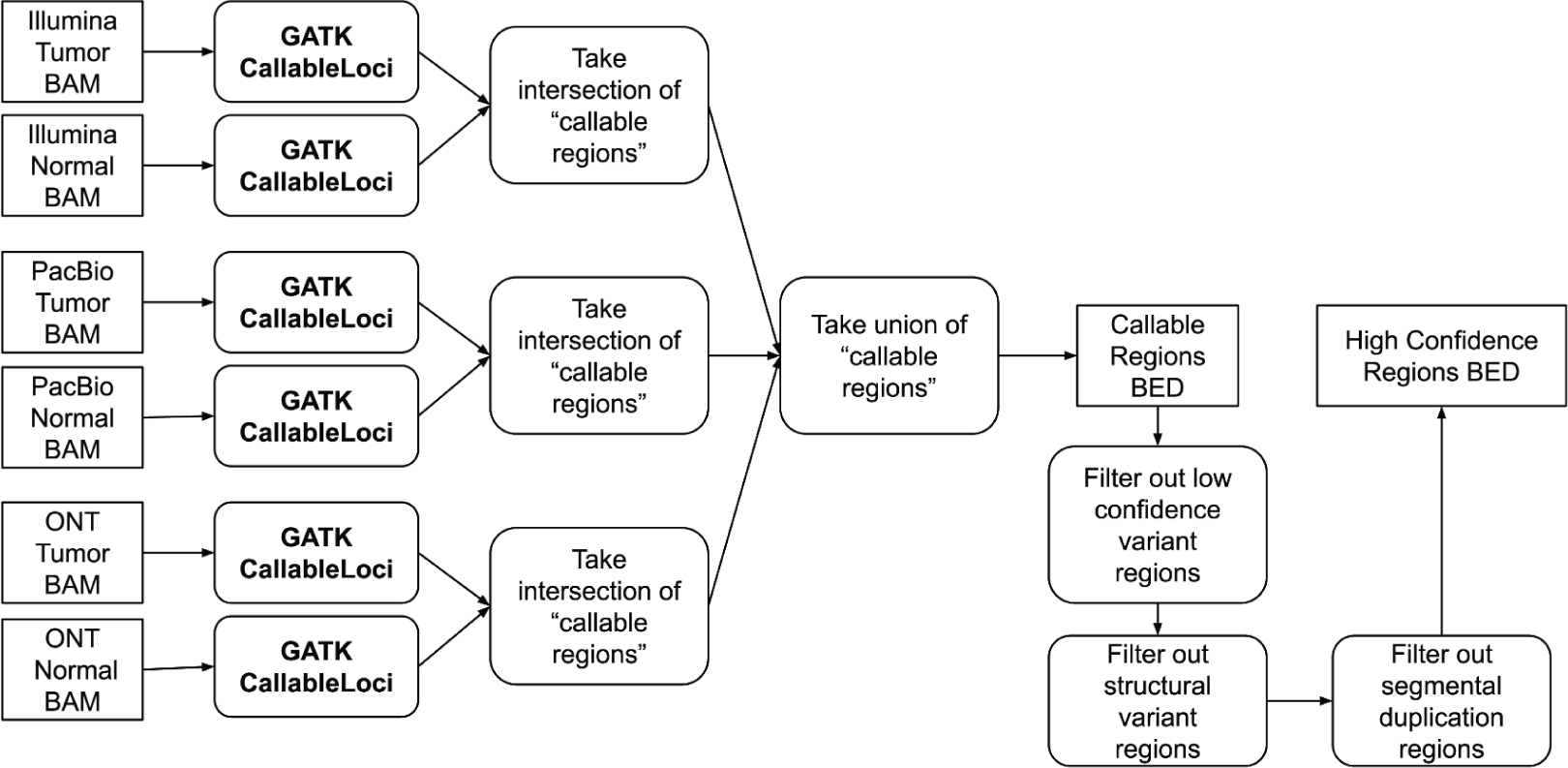
Steps for generating high confidence regions BED files.

**Supplementary Figure 2:**
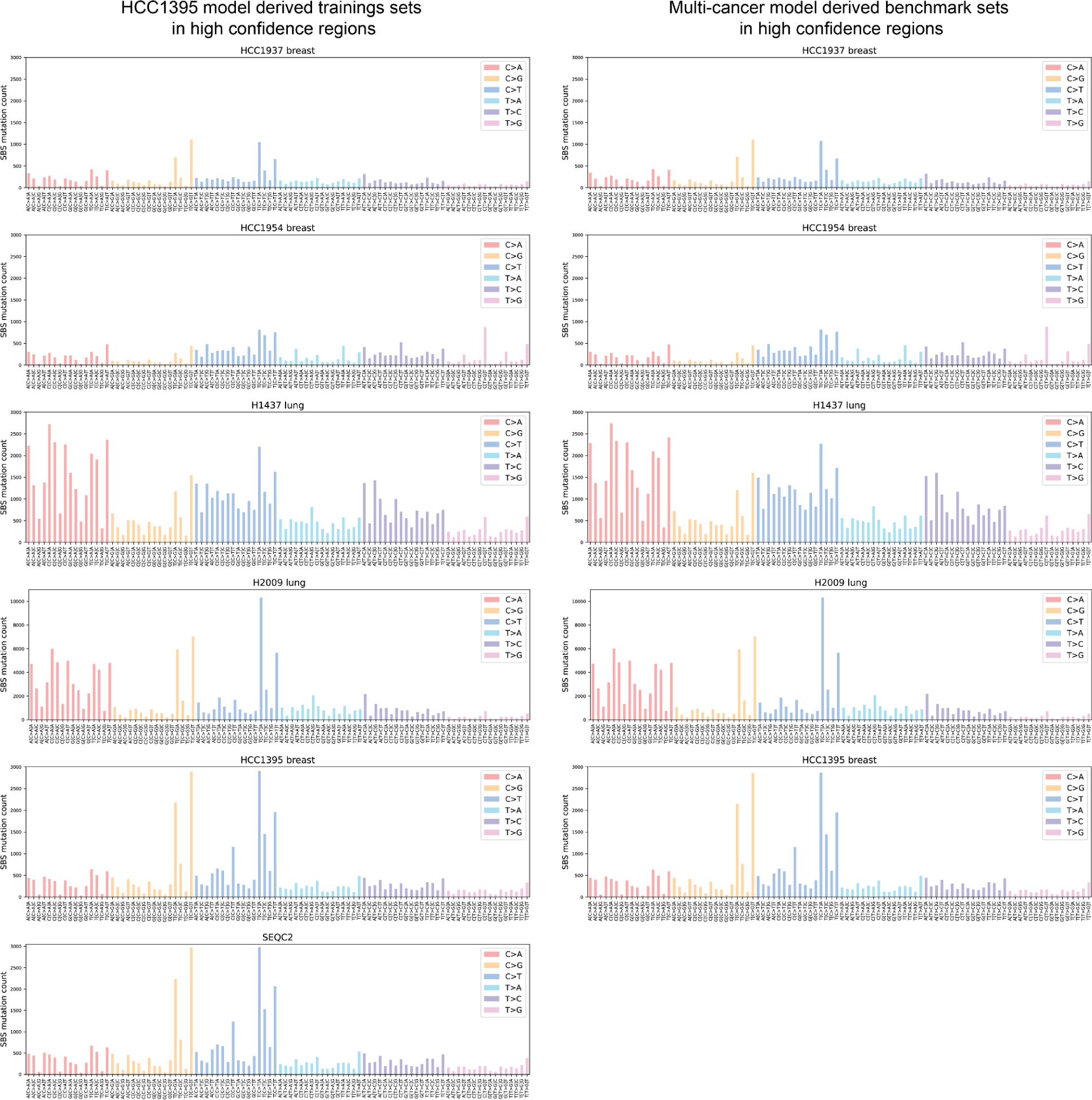
SBS-96 counts for SEQC2 benchmark and each benchmark set derived from DeepSomatic HCC1395 models (left) and multi-cancer models (right) in high-confidence regions.

**Supplementary Figure 3:**
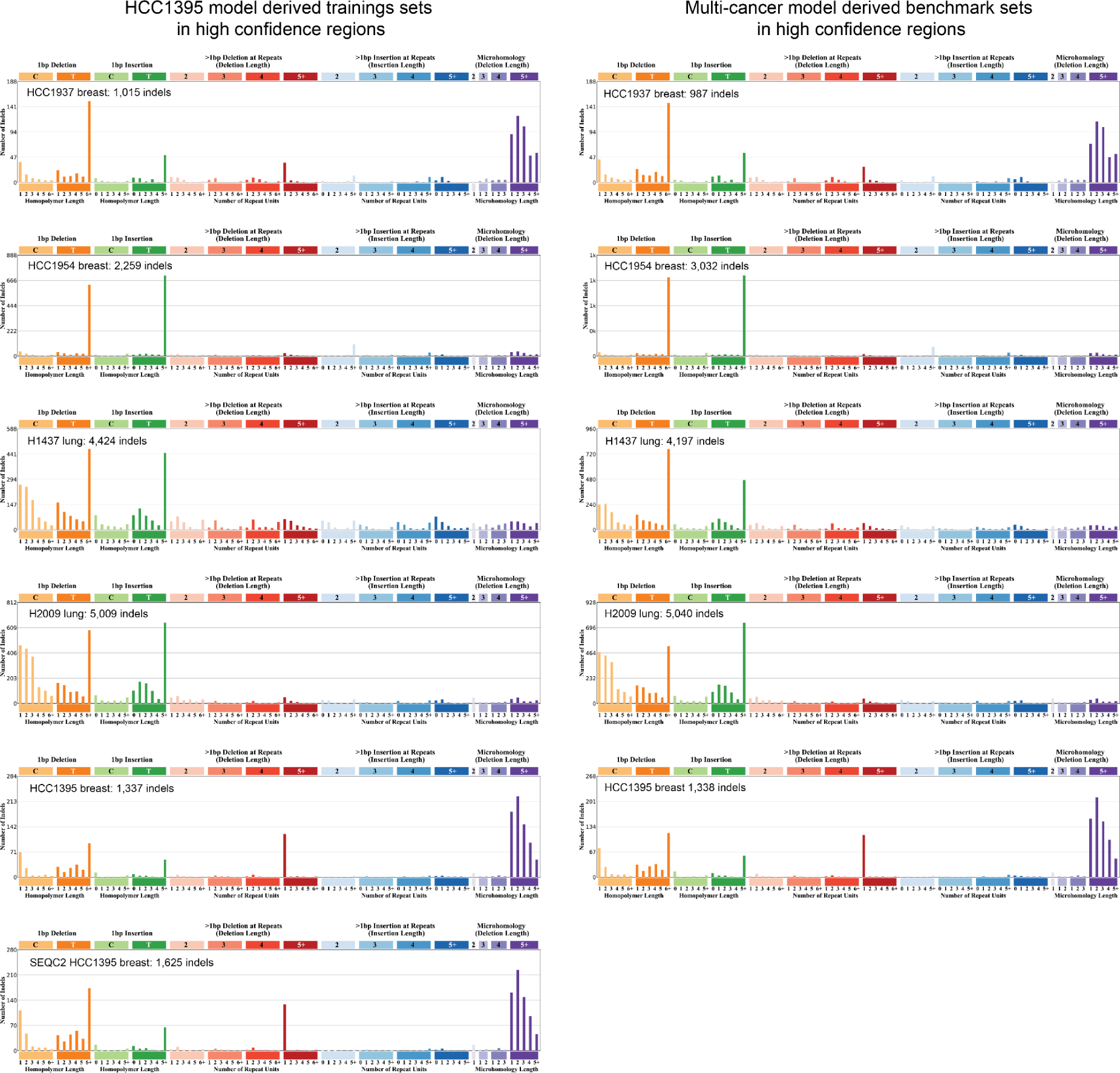
indel classification counts for SEQC2 benchmark and each benchmark set derived from DeepSomatic HCC1395 models **(left)** and multi-cancer models **(right)** in high confidence regions. Plots were generated using SigProfileMatrixGenerator.

**Supplementary Figure 4:**
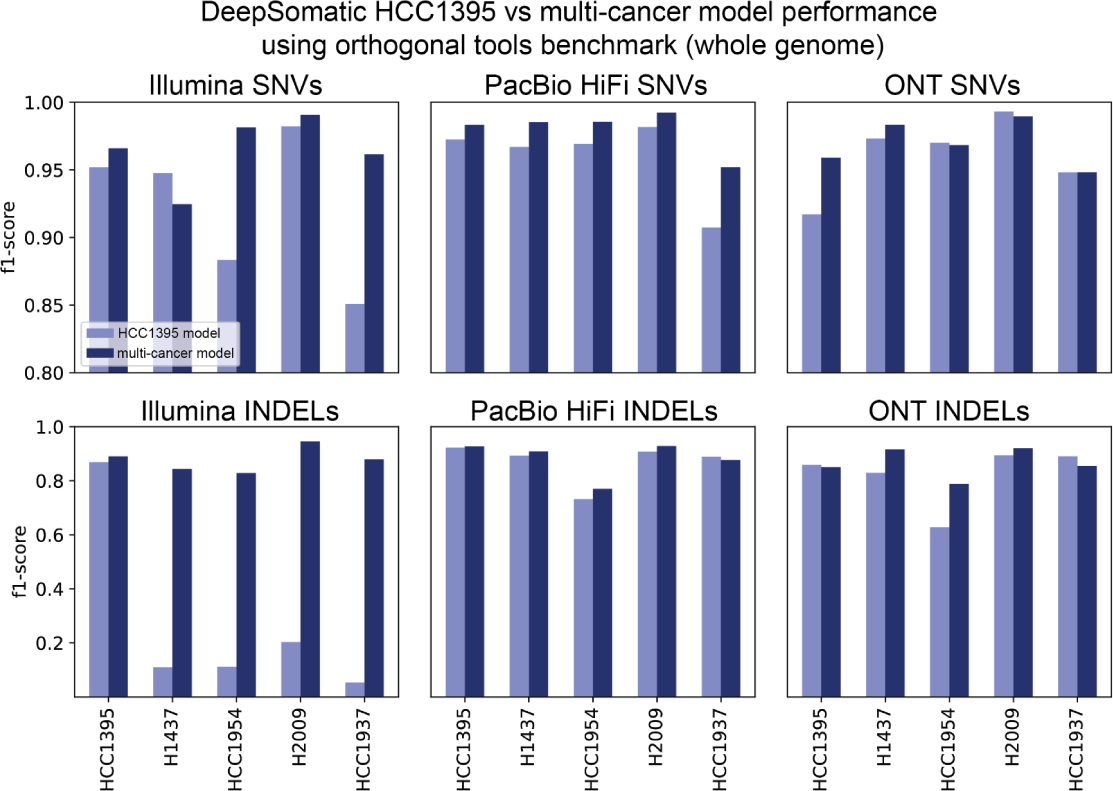
Performance of DeepSomatic HCC1395 model vs multi-cancer model against orthogonal tools benchmark on the whole genome.

**Supplementary Figure 5:**
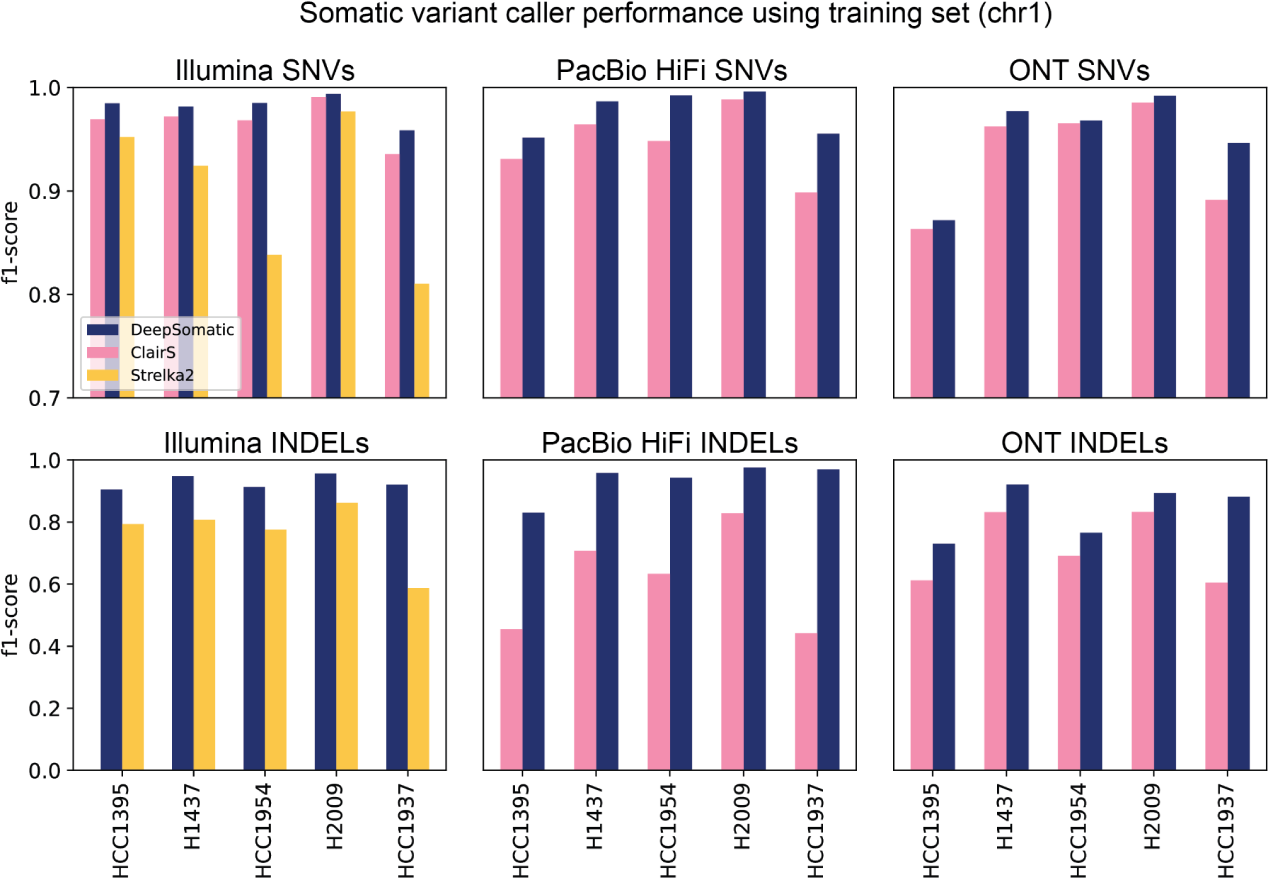
Performance of somatic variant callers against the training set derived from DeepSomatic HCC1395 models.

**Supplementary Figure 6:**
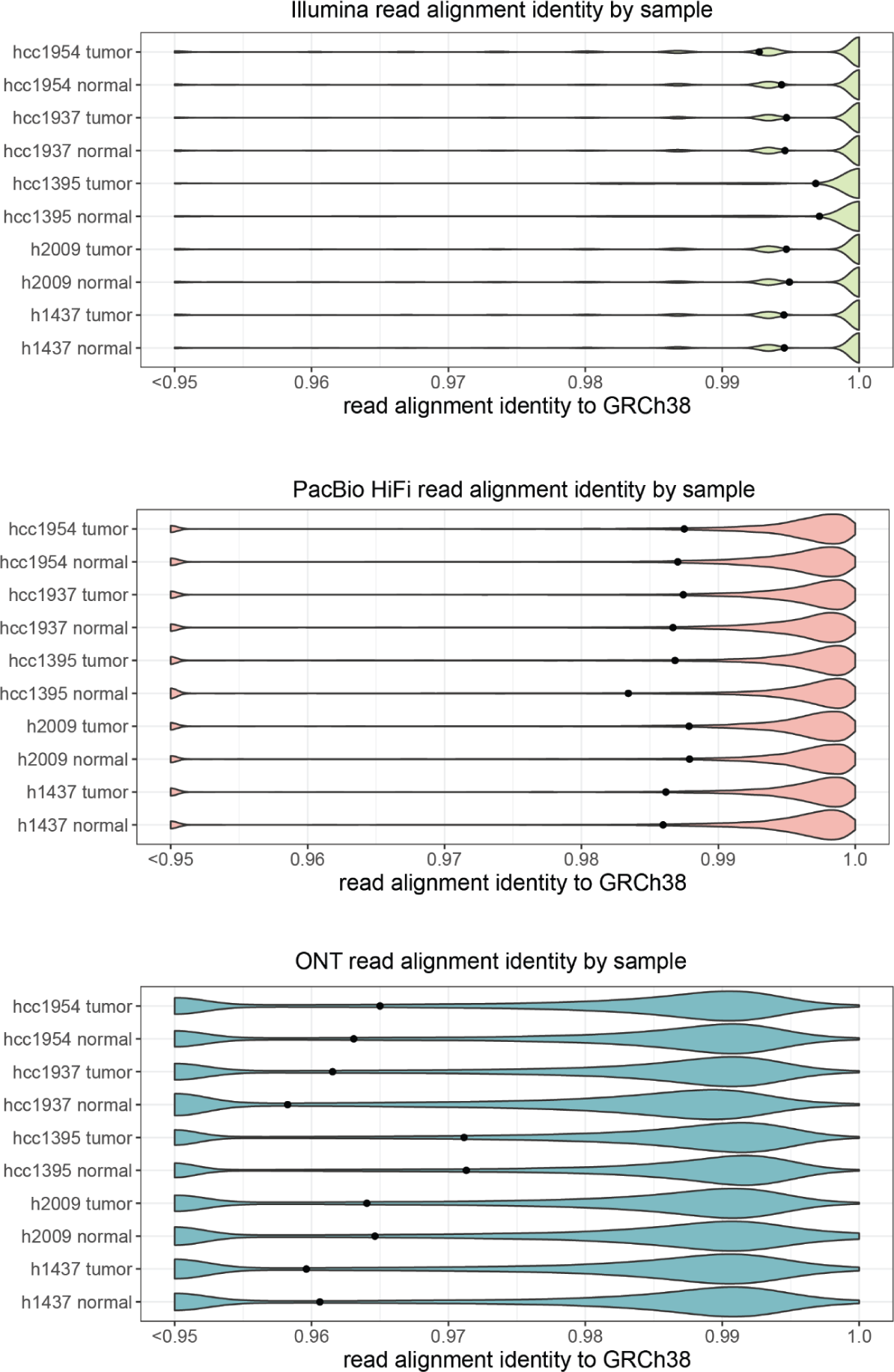
Read alignment identity to GRCh38 by sequencing technology and cell line sample. Points represent the mean for each cell line sample.

**Supplementary Figure 7:**
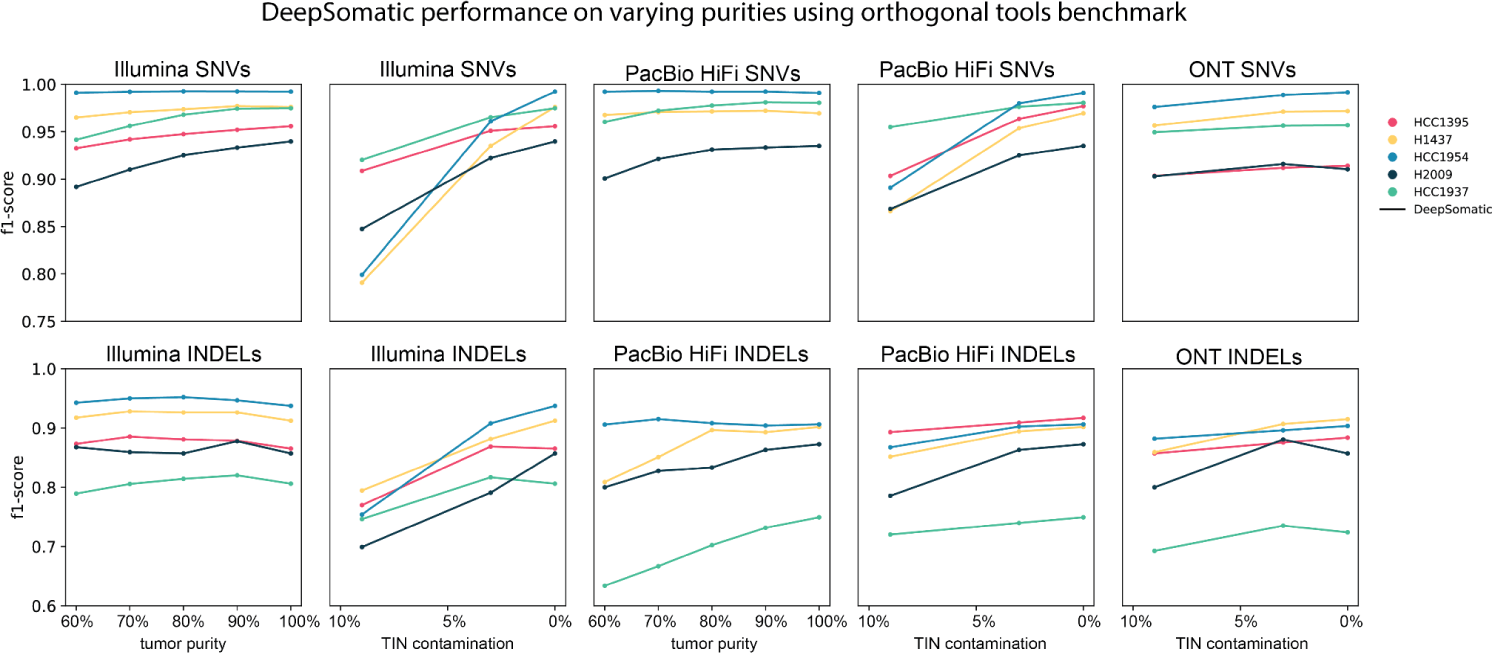
Performance of DeepSomatic multi-cancer model on various tumor purities and TIN contamination against orthogonal tools benchmark on chromosome 1.

**Supplementary Figure 8:**
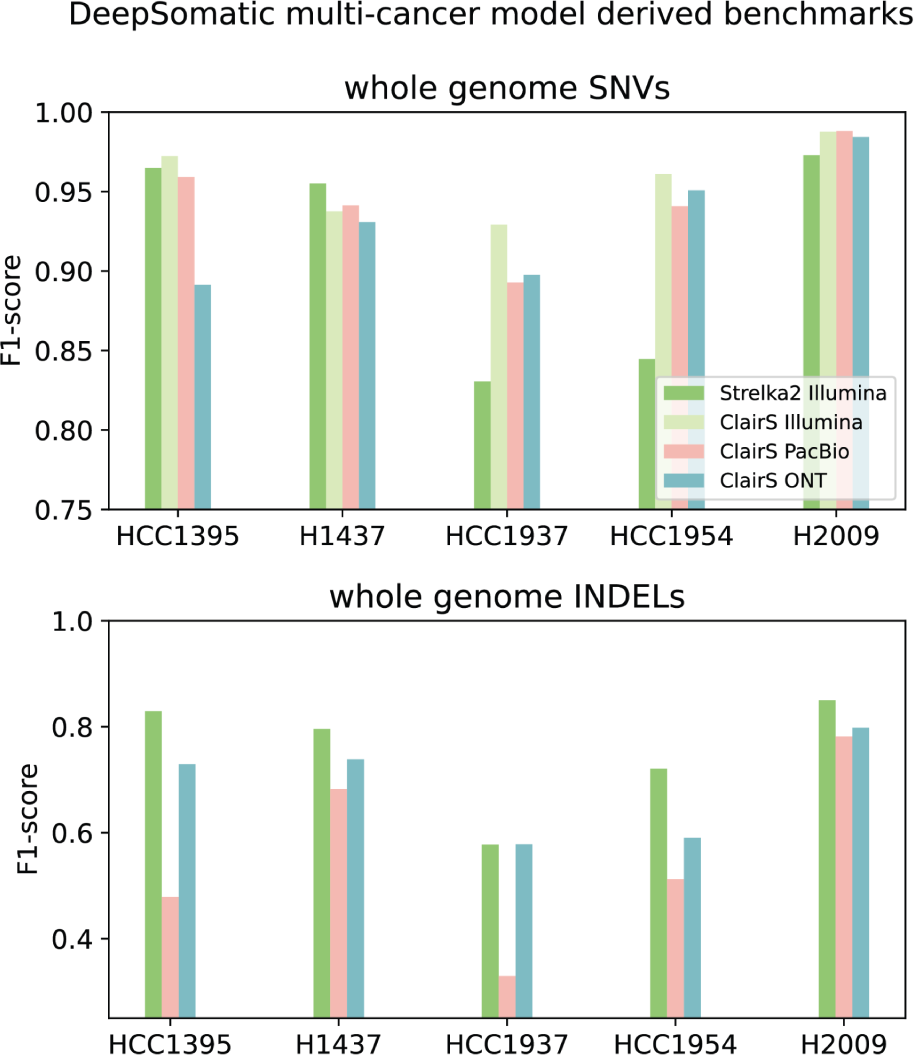
Benchmarking somatic variant callers using DeepSomatic multi-cancer model derived benchmarks.

**Supplementary Figure 9:**
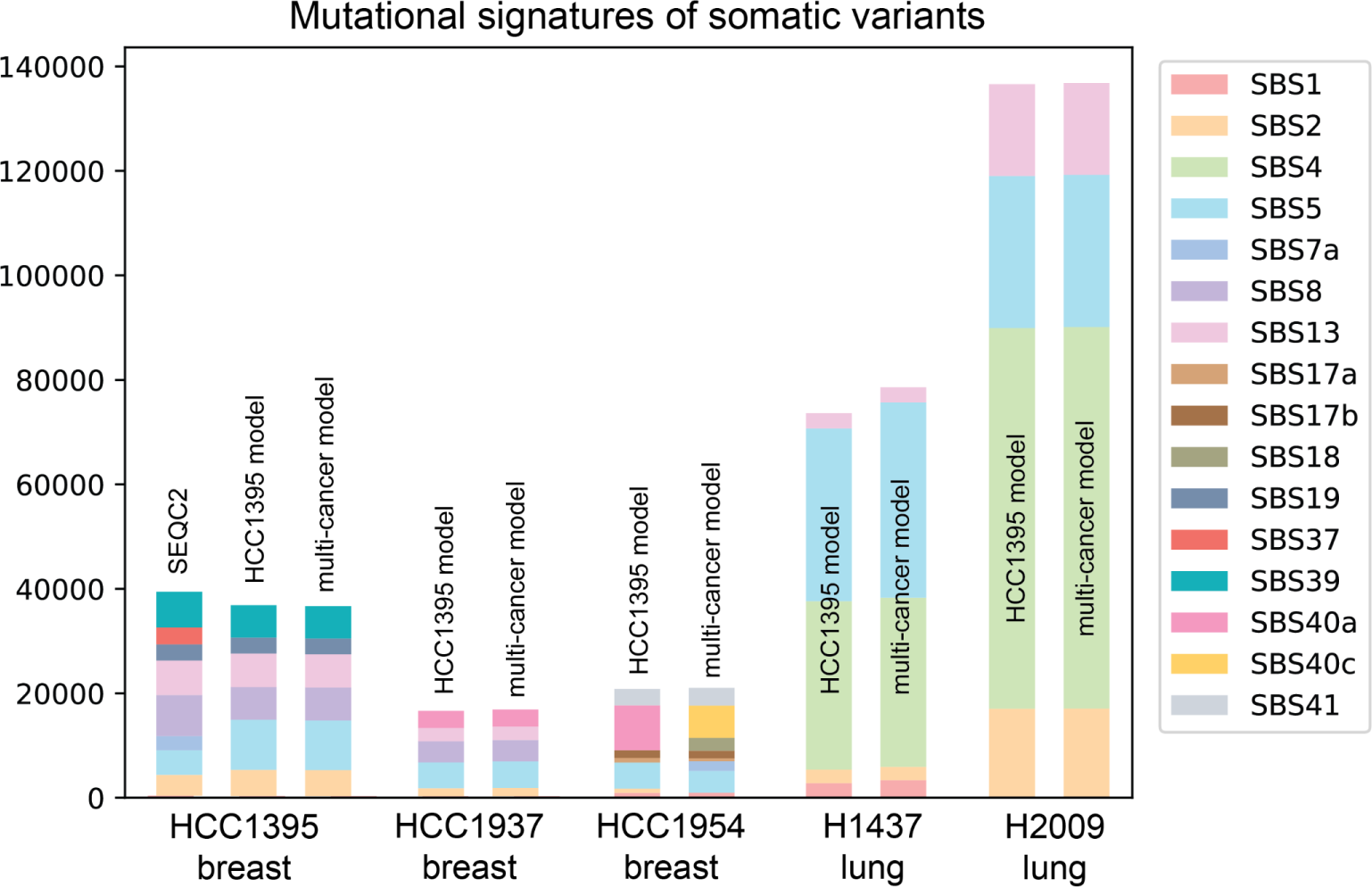
Comparison of mutational signatures of somatic variants called by DeepSomatic HCC1395 models and multi-cancer models as well as the SEQC2 benchmark, showing them to be highly consistent.

**Supplementary Figure 10:**
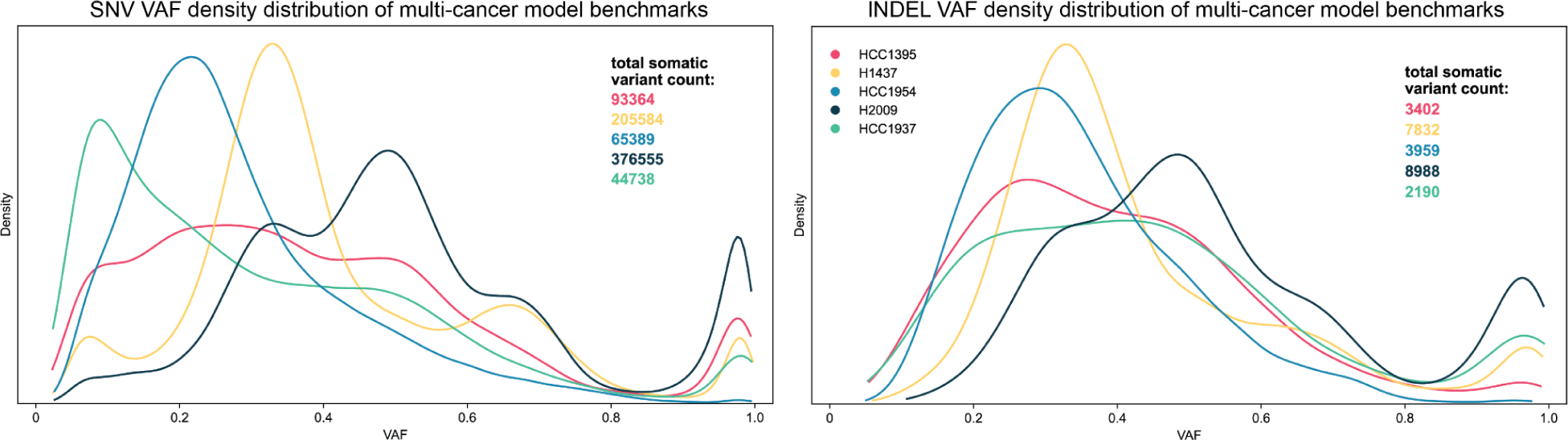
Variant allele frequency (VAF) distributions of somatic variant sets derived using DeepSomatic multi-cancer models in high confidence regions, represented as Kernel Density Estimate (KDE) plots.

**Figure 11:**
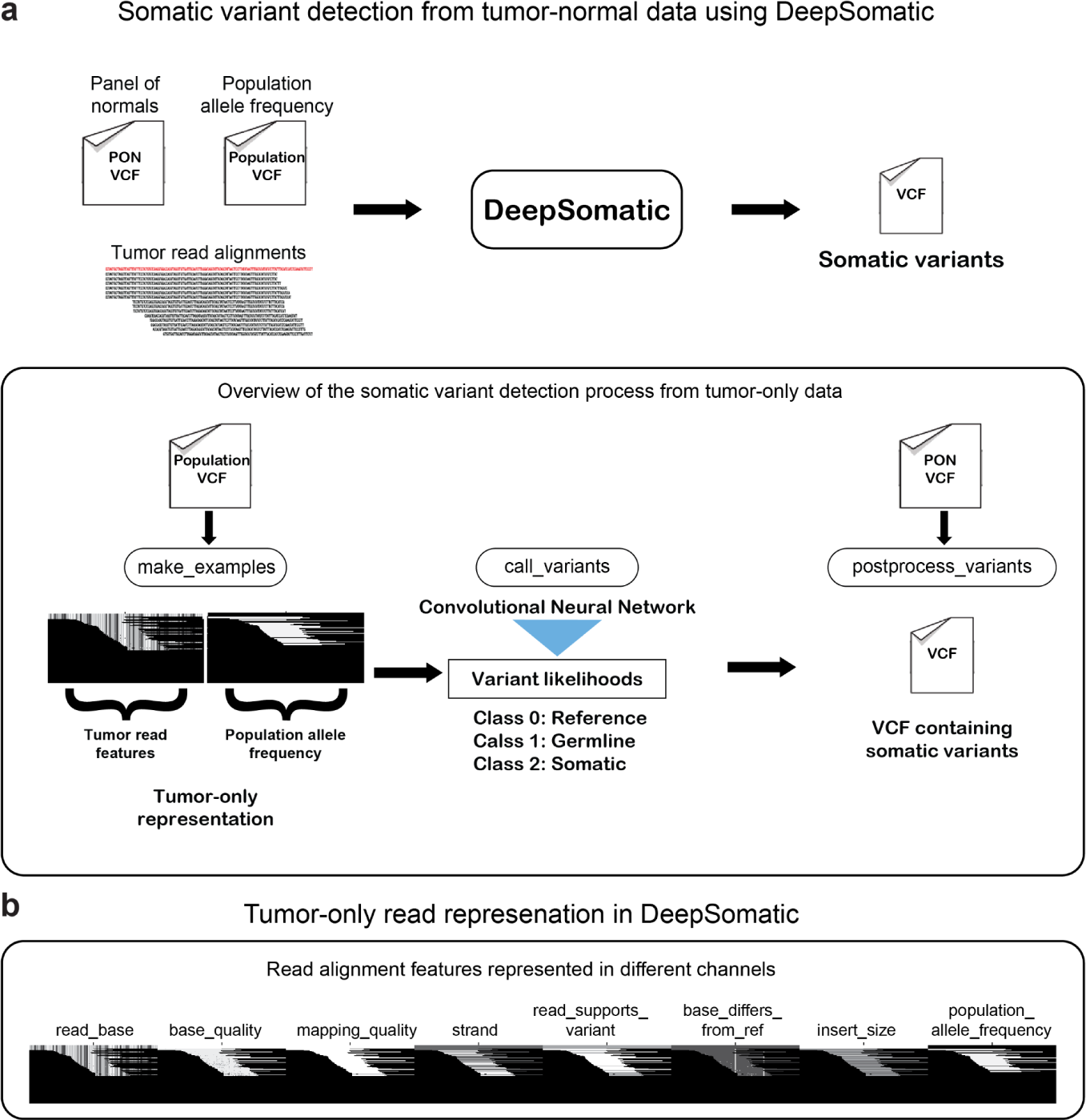
DeepSomatic tumor-only variant calling overview. **(a)** Overview of DeepSomatic tumor-only mode. **(b)** Tumor-only read alignment features represented including population allele frequency.

## Command lines

### Read Alignment

Minimap2 (v2.26)

https://github.com/lh3/minimap2

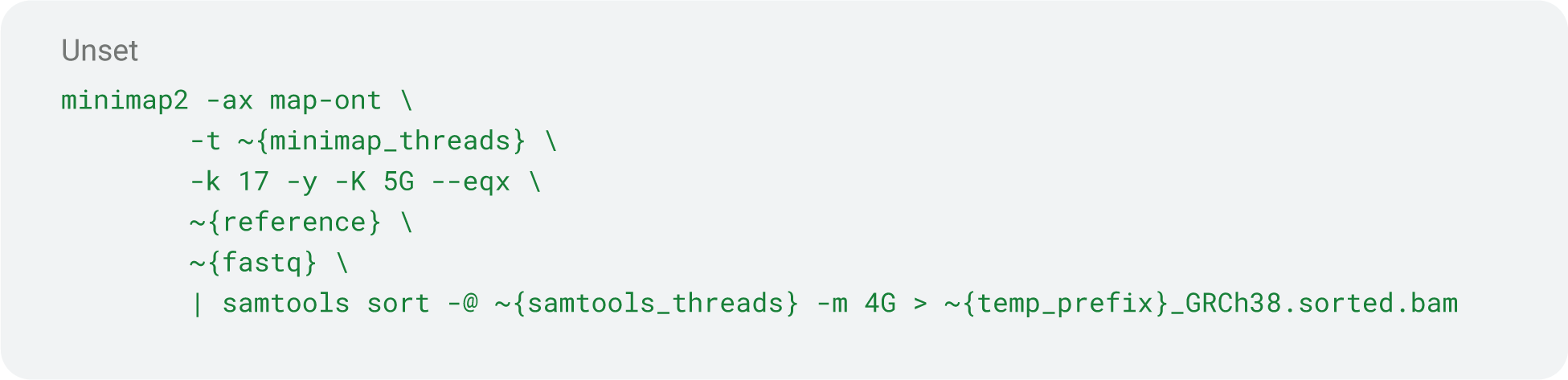

Pbmm2 (docker quay.io/biocontainers/pbmm2:1.13.1--h9ee0642_0)

https://github.com/PacificBiosciences/pbmm2

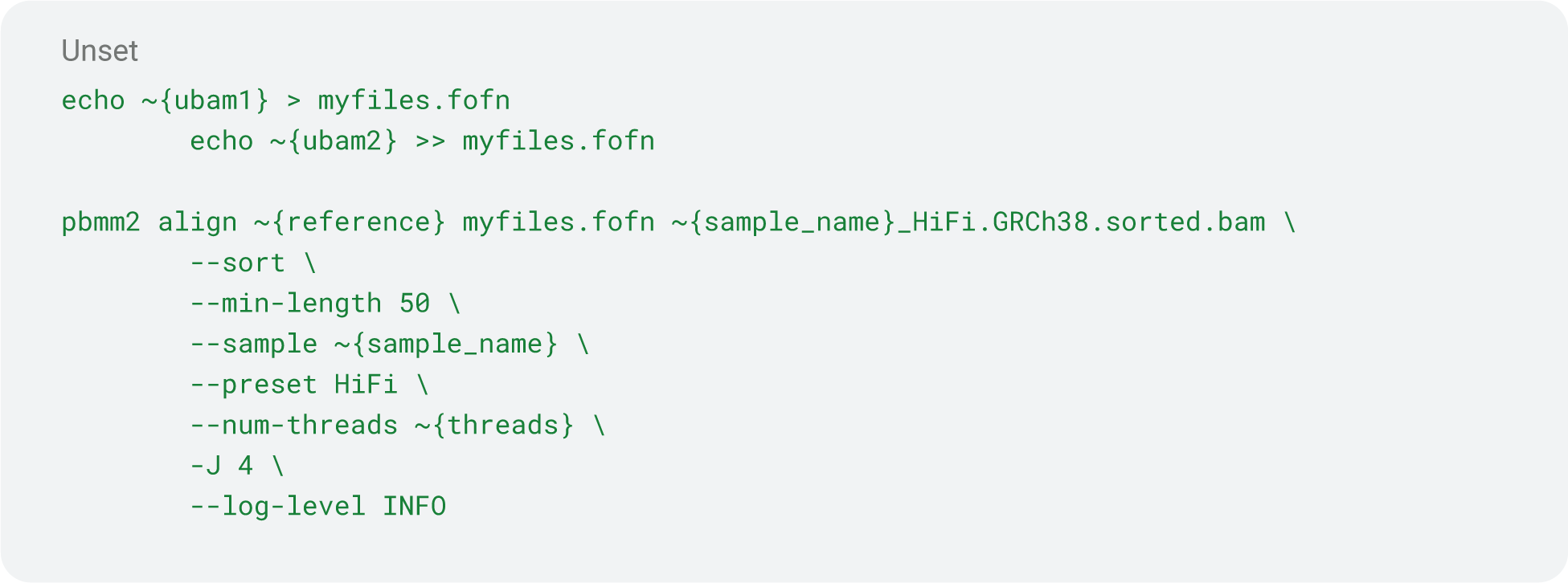

Bwa-mem2 (v2.2.1)

https://github.com/bwa-mem2/bwa-mem2

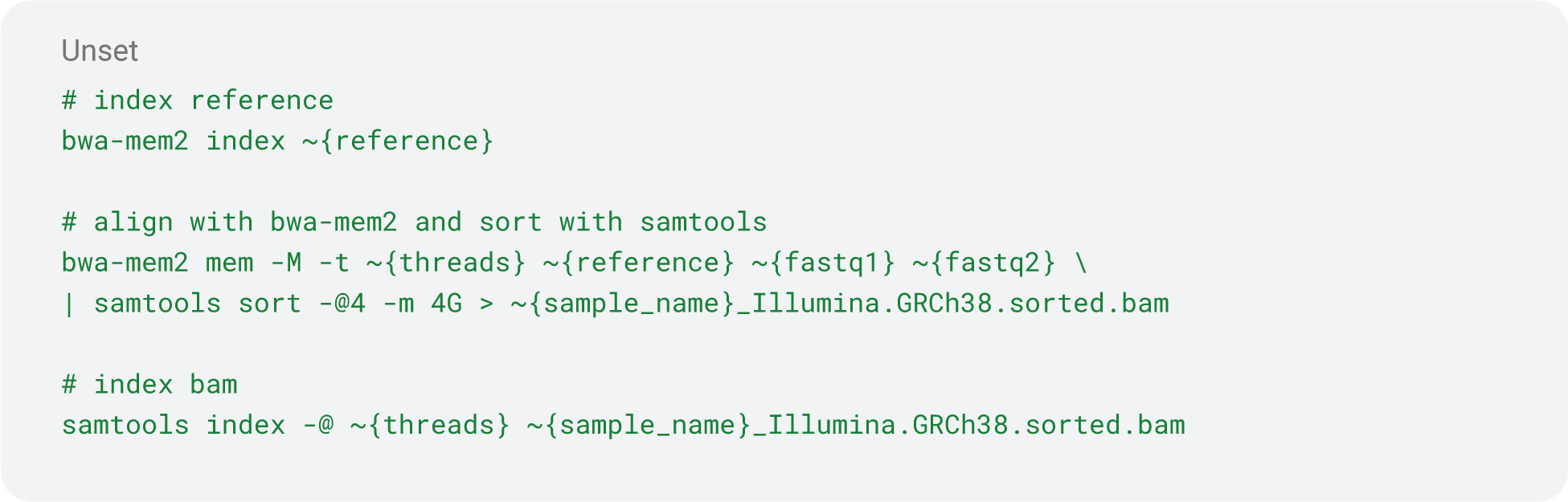

Calculating depth of BAMs (samtools v1.13)

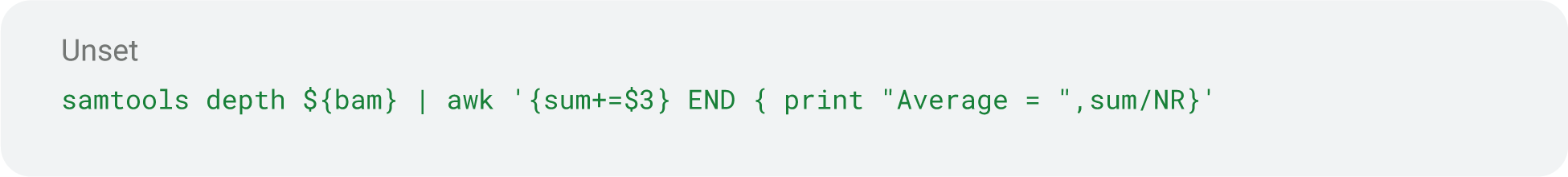

Calculating alignment identity and N50 (wambam)

https://github.com/nanoporegenomics/wambam

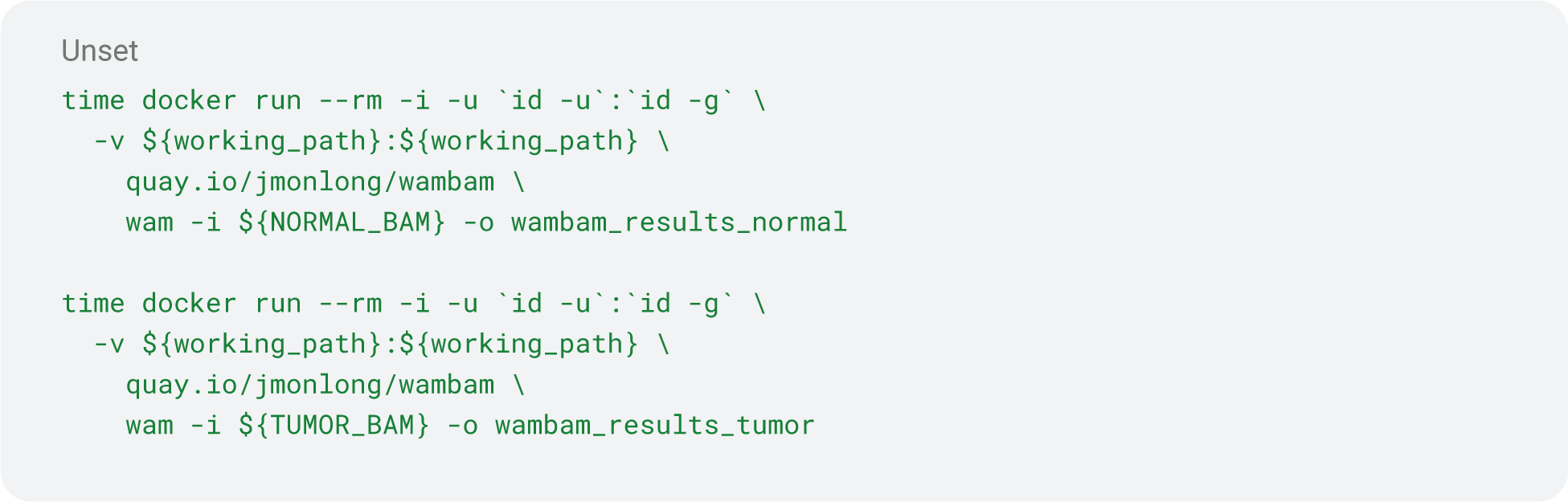

### Variant Calling

DeepSomatic

https://github.com/google/deepsomatic

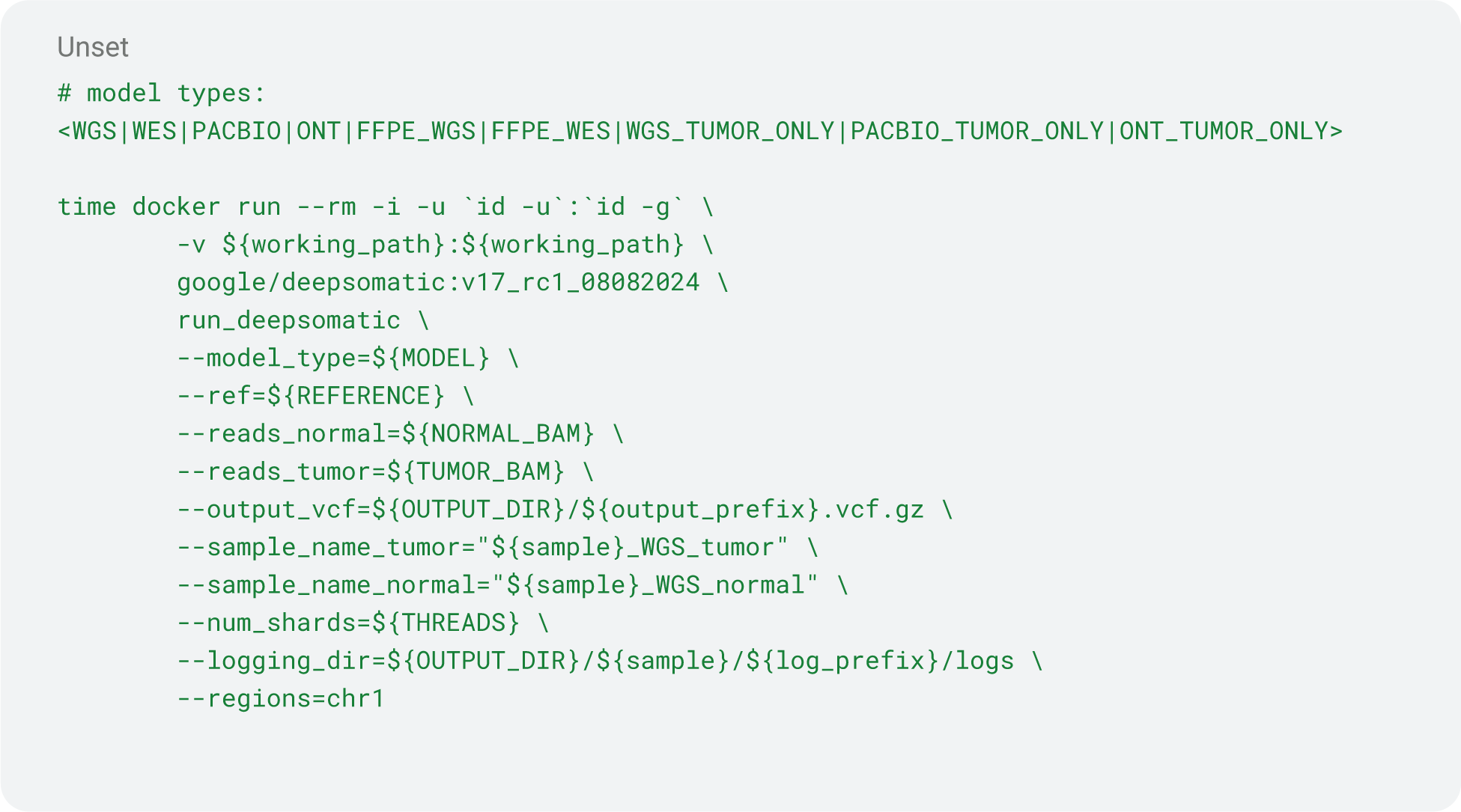

ClairS (v0.2.0)

https://github.com/HKU-BAL/ClairS

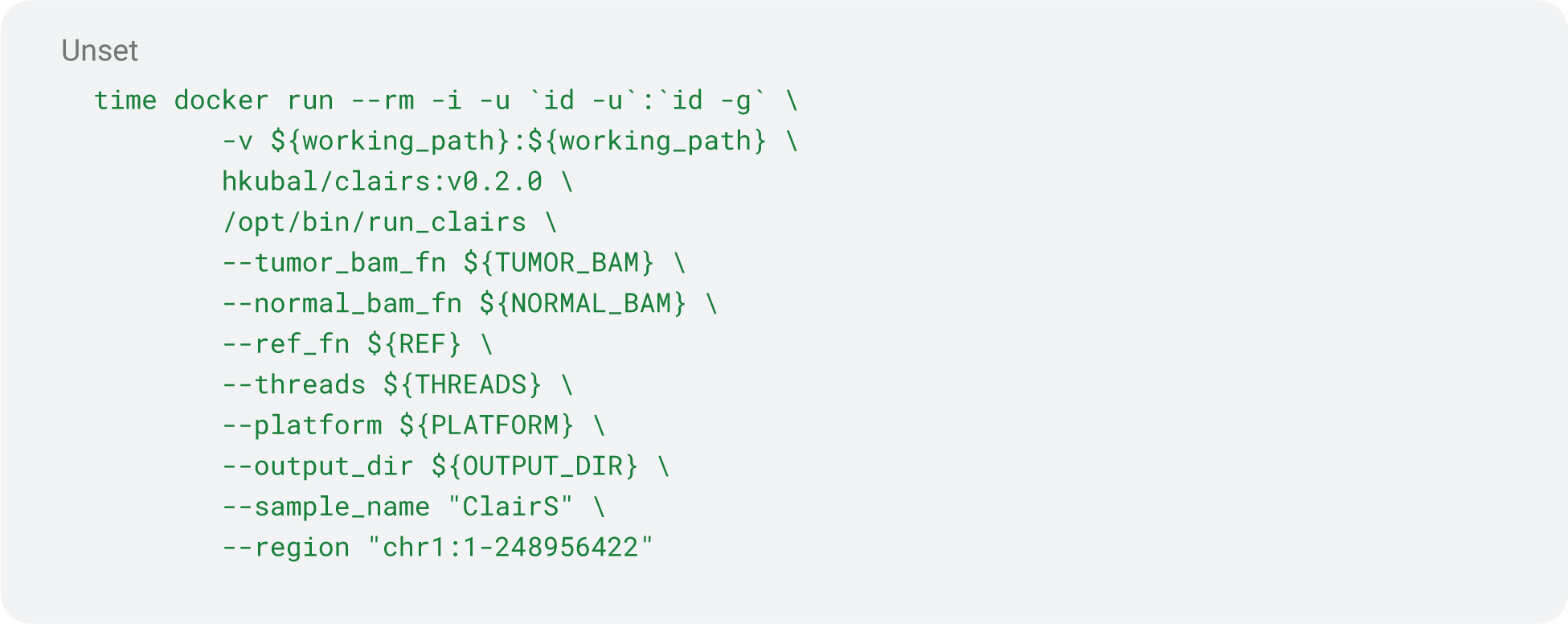

Strelka2 (v2.9.10)

https://github.com/Illumina/strelka

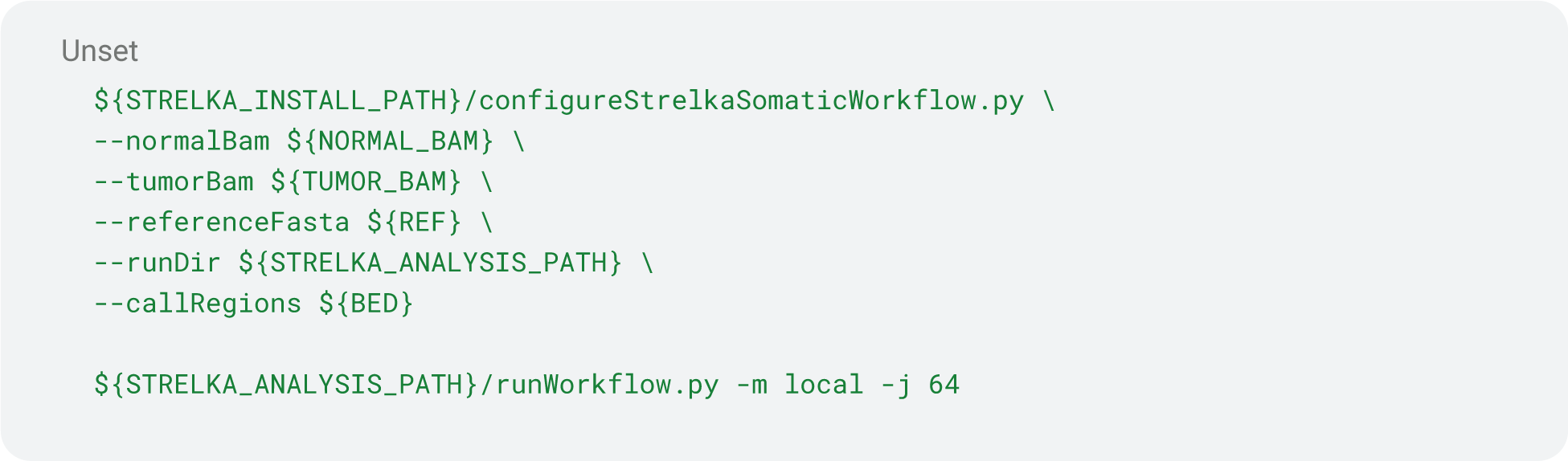

DeepVariant HYBRID_PACBIO_ILLUMINA (v1.6.1)

https://github.com/google/deepvariant

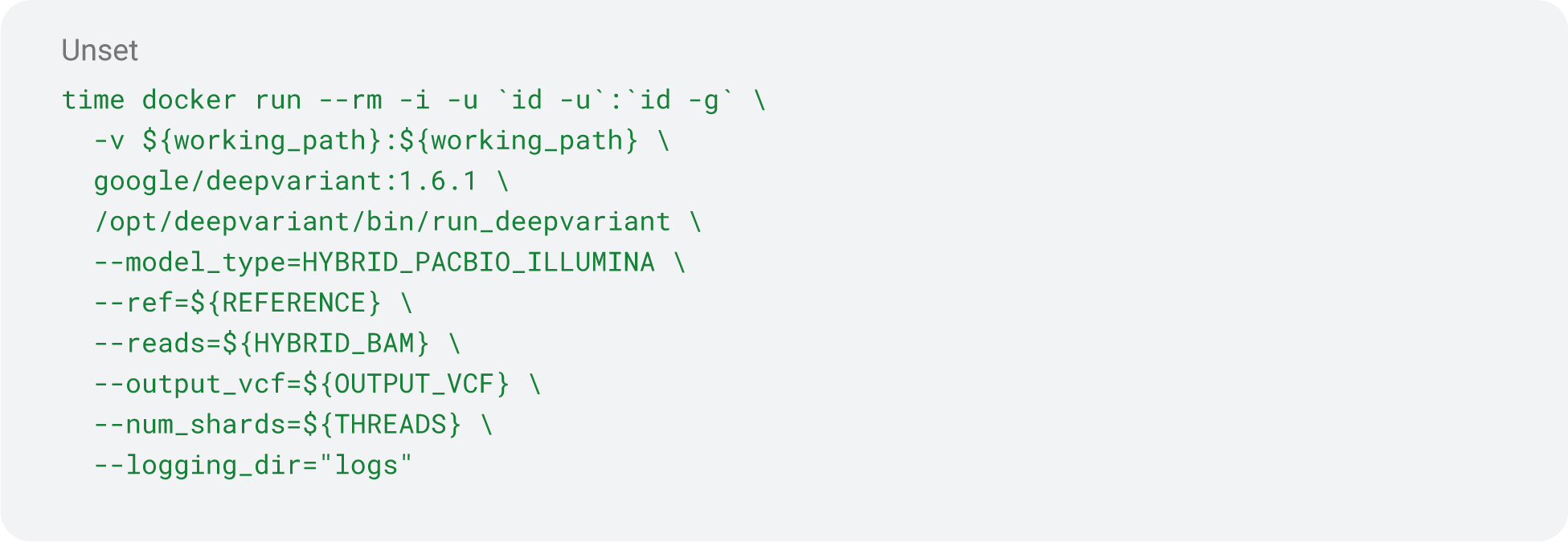

Generating training sets

https://github.com/jimin001/DeepSomatic_manuscript/blob/main/vcf_intersection_complex_v2.py

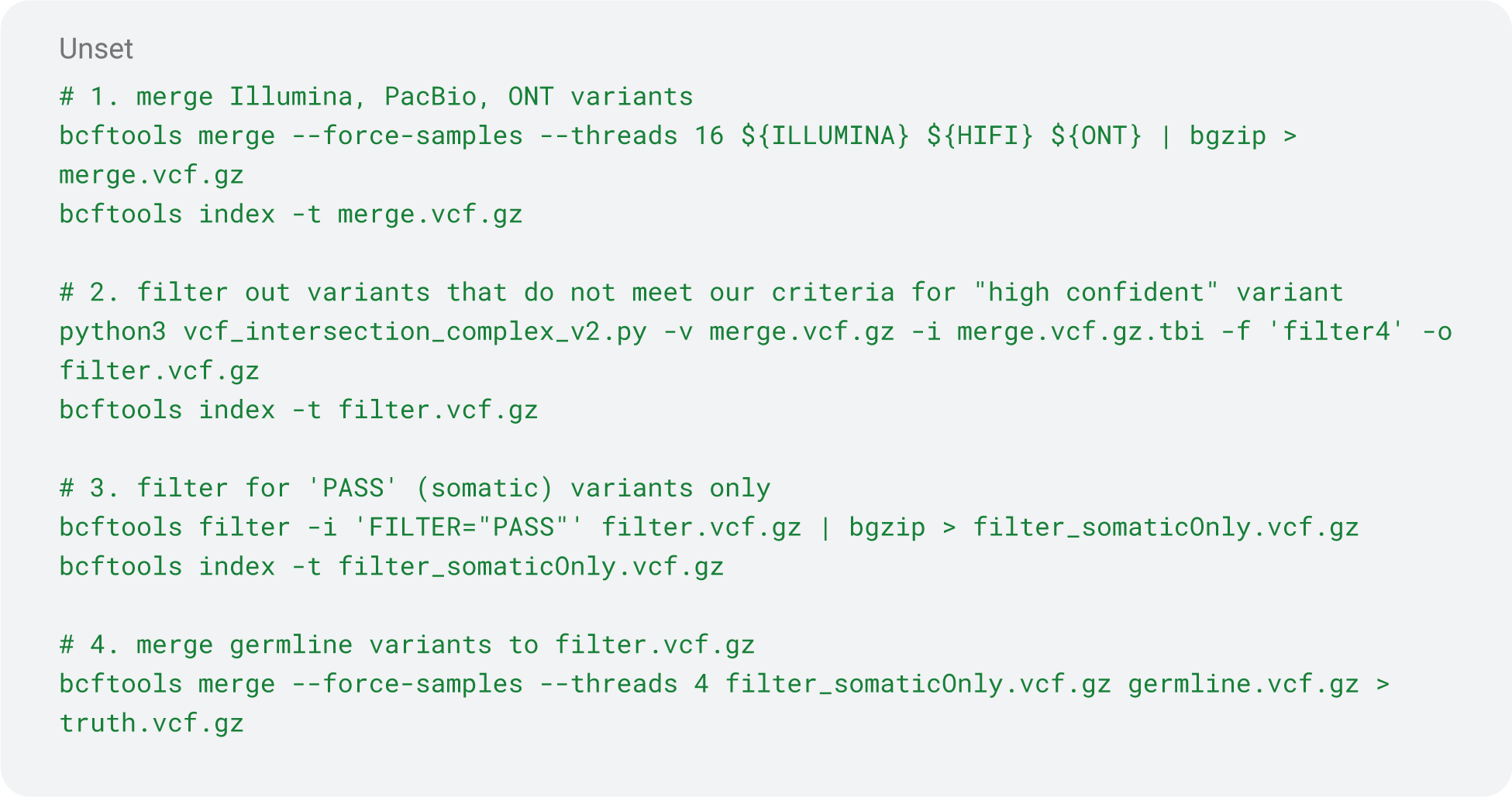

Generating high confidence region BED files

https://github.com/jimin001/DeepSomatic_manuscript/blob/main/vcf_to_bed_v4.py

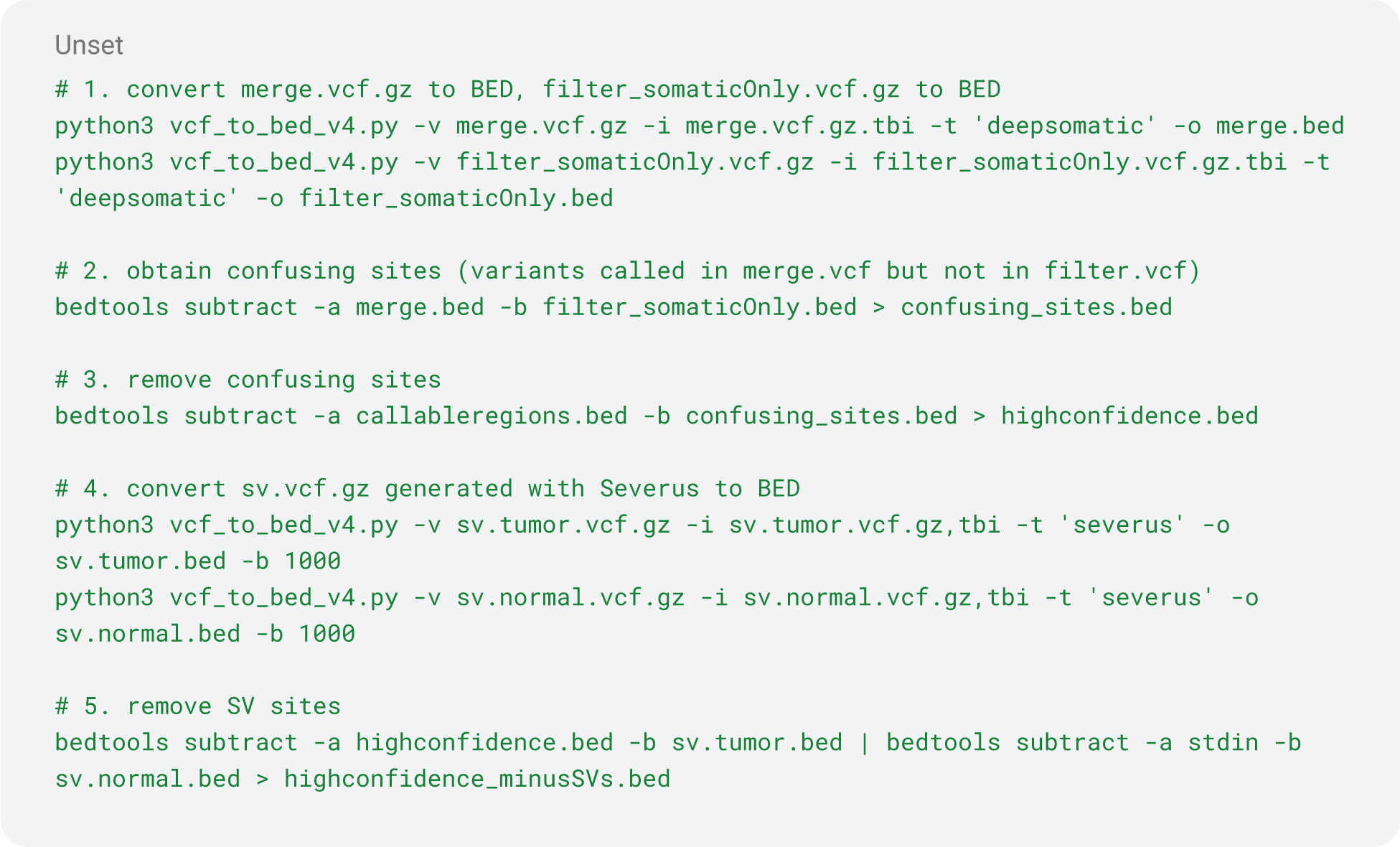

Generating callable regions

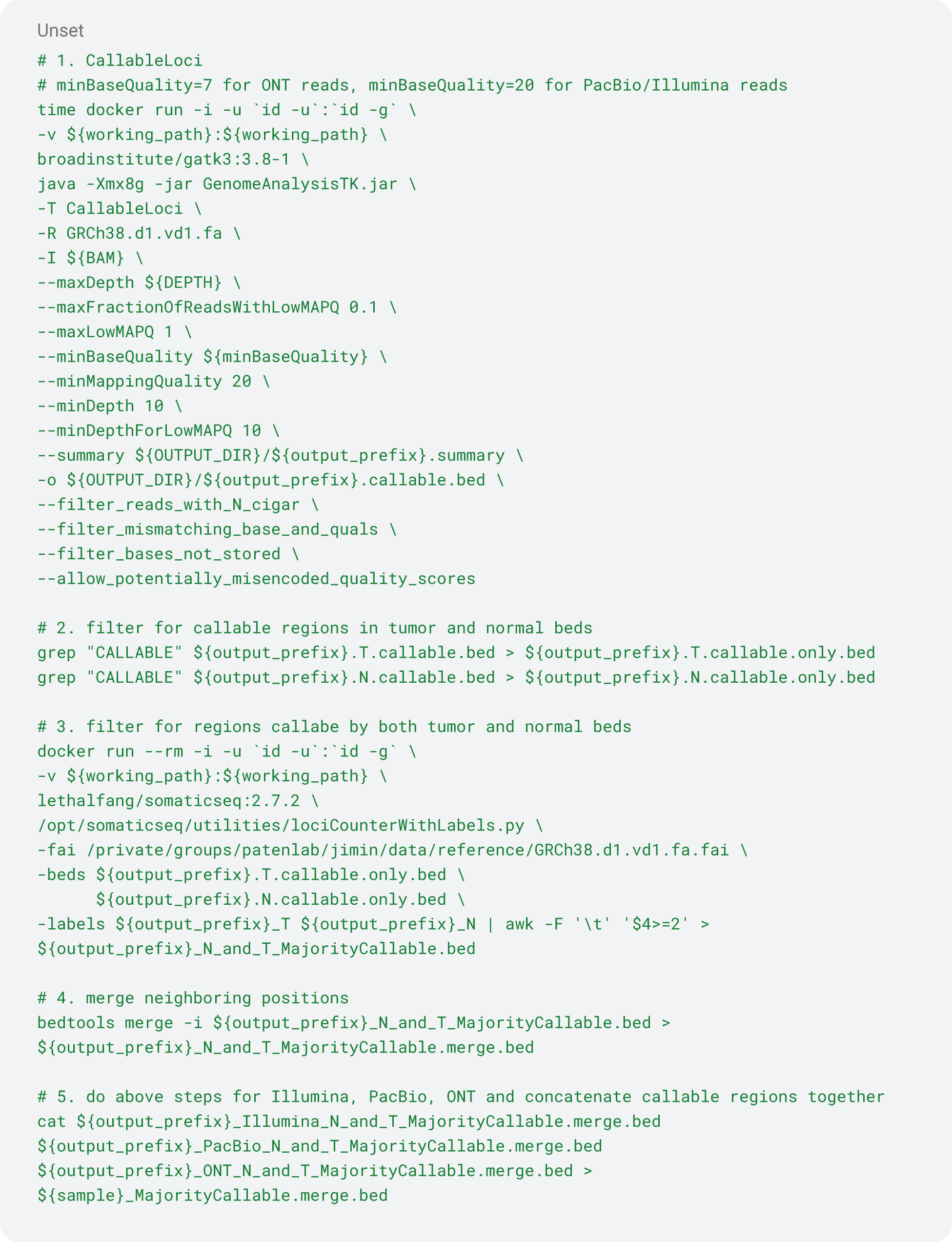

Generating titration bams

https://github.com/jimin001/DeepSomatic_manuscript/blob/main/split_bam_tumor.sh

https://github.com/jimin001/DeepSomatic_manuscript/blob/main/split_bam_normal.sh

https://github.com/jimin001/DeepSomatic_manuscript/blob/main/tumor_purity_titration.sh

https://github.com/jimin001/DeepSomatic_manuscript/blob/main/normal_purity_titration.sh

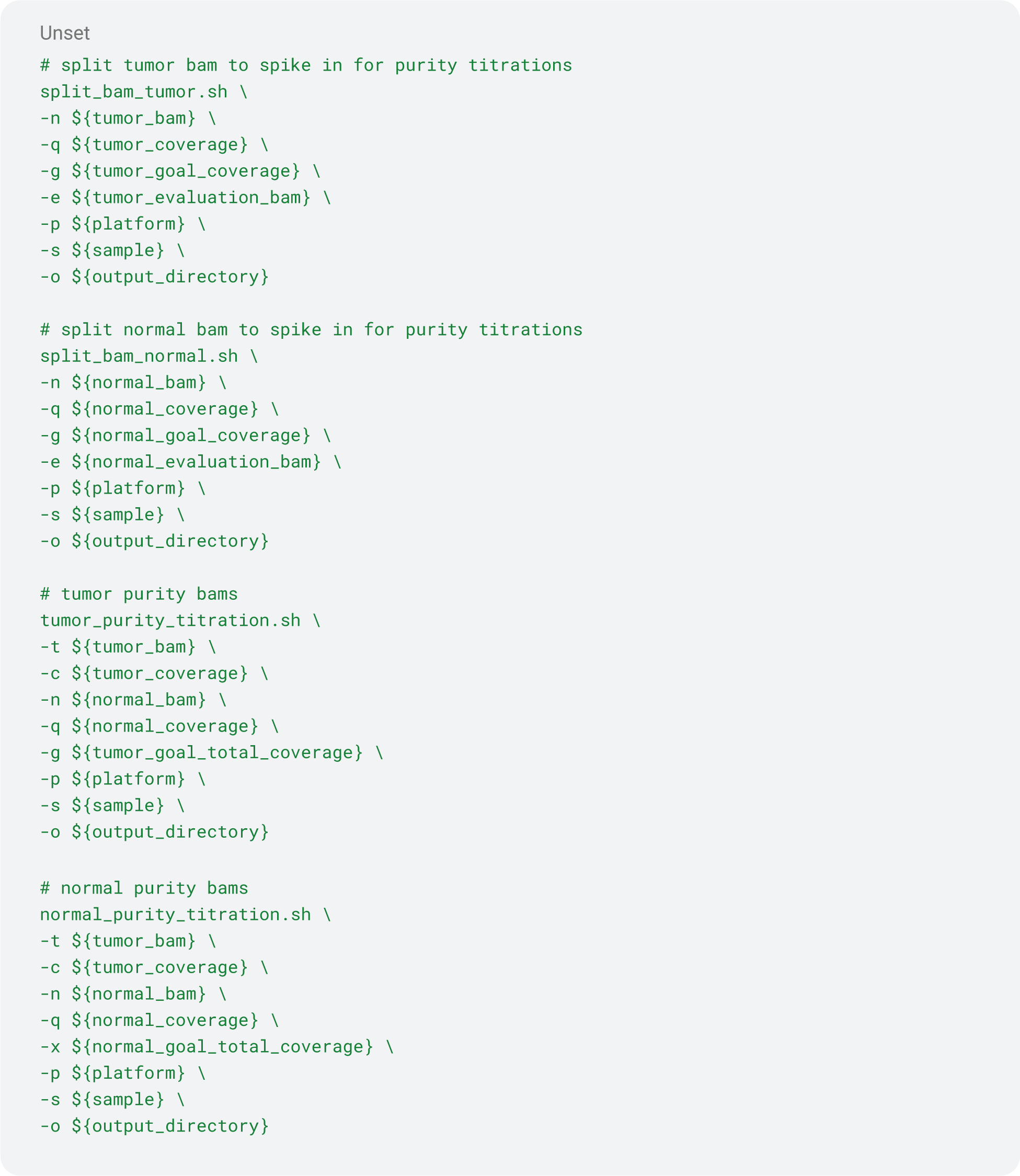

Mutational signature analysis

https://github.com/AlexandrovLab/SigProfilerAssignment

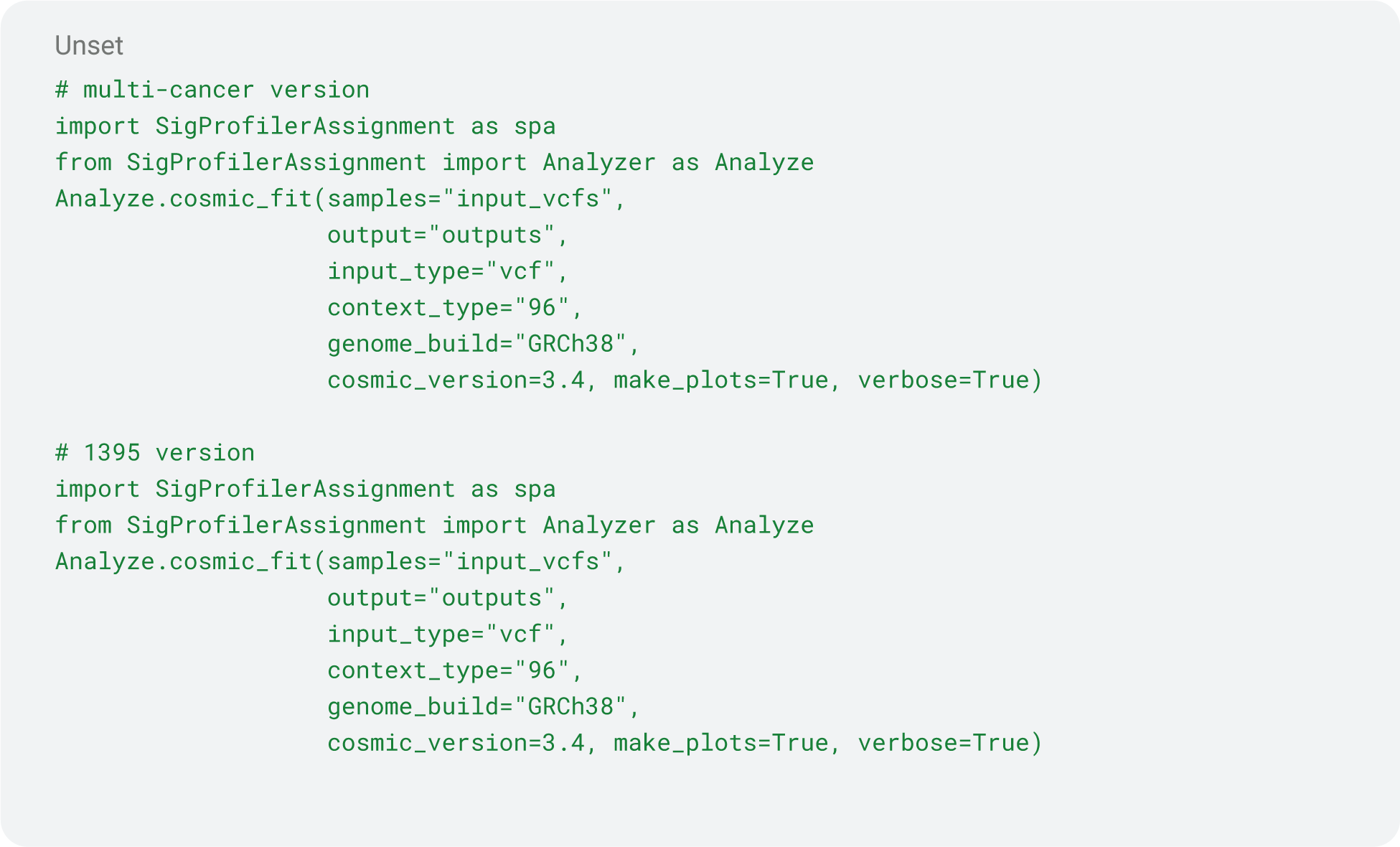

Mutational matrix generation

https://github.com/AlexandrovLab/SigProfilerMatrixGenerator

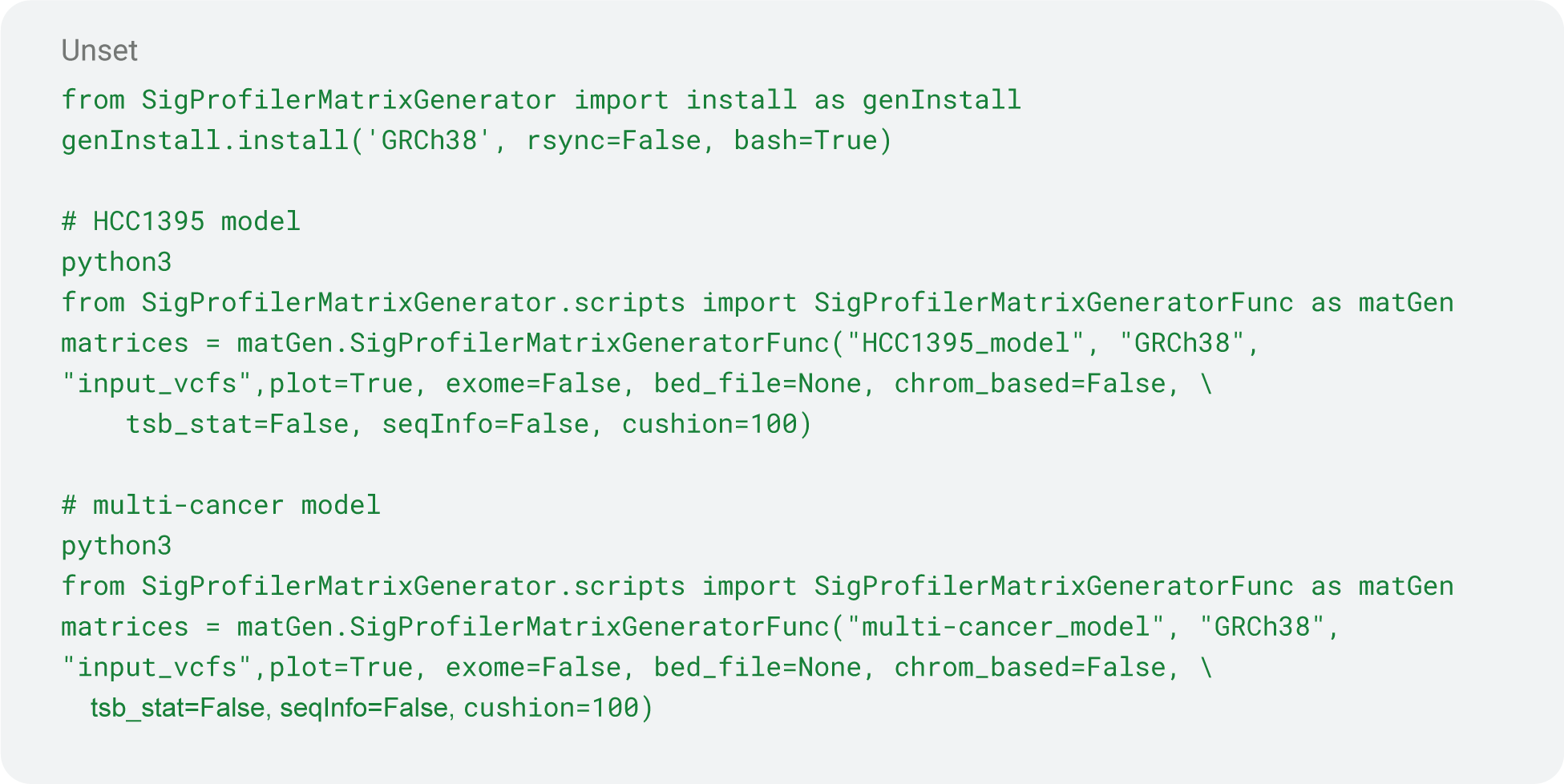

### Benchmarking

Som.py

https://github.com/Illumina/hap.py/blob/master/doc/sompy.md

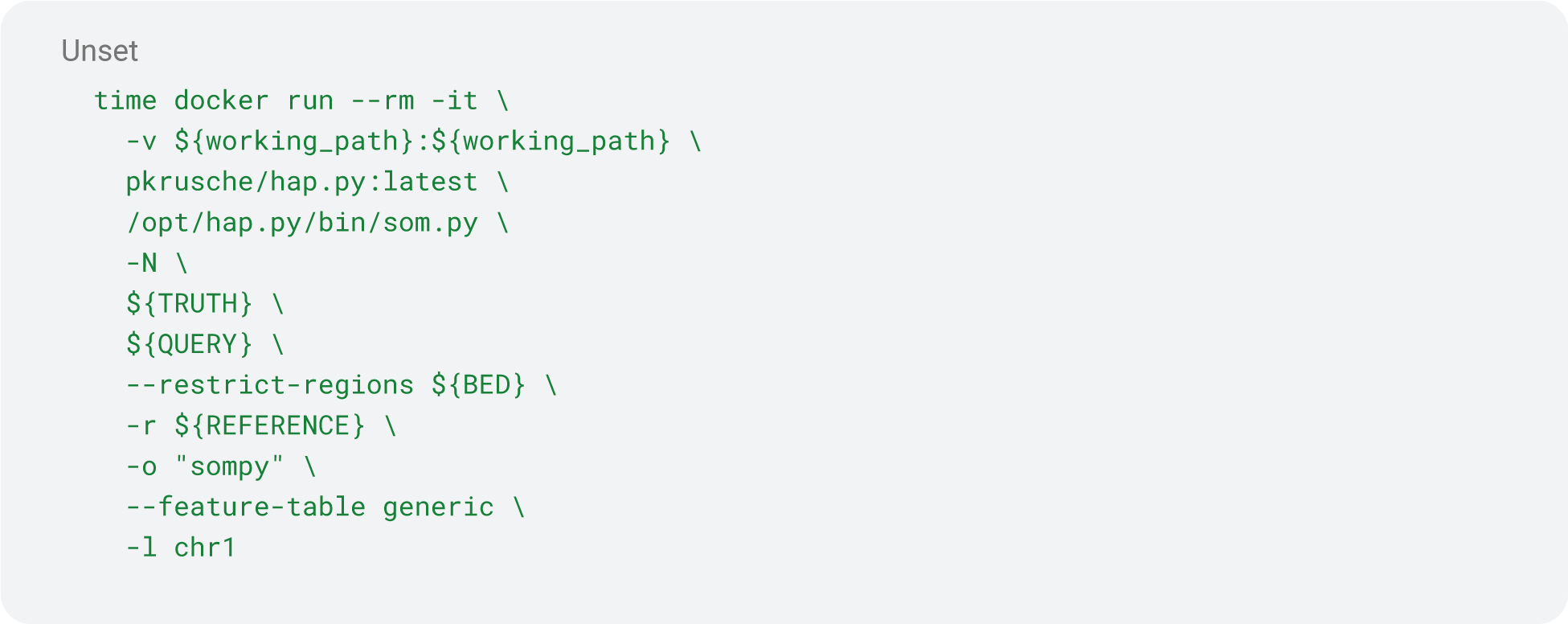

